# Metagenome-assembled genomes of phytoplankton communities across the Arctic Circle

**DOI:** 10.1101/2020.06.16.154583

**Authors:** A. Duncan, K. Barry, C. Daum, E. Eloe-Fadrosh, S. Roux, S. G. Tringe, K. Schmidt, K. U. Valentin, N. Varghese, I. V. Grigoriev, R. Leggett, V. Moulton, T. Mock

**Affiliations:** School of Computing Sciences, University of East Anglia, Norwich Research Park, NR47TJ, Norwich, U.K.; DOE-Joint Genome Institute, 1 Cyclotron Road, Berkeley, CA 94720, U.S.A.; School of Environmental Sciences, University of East Anglia, Norwich Research Park, NR47TJ, Norwich, U.K.; Alfred-Wegener Institute for Polar and Marine Research, Am Handelshafen 12, 27570 Bremerhaven, Germany; Earlham Institute, Norwich Research Park, Norwich, NR4 7UG, U.K.

## Abstract

Phytoplankton communities significantly contribute to global biogeochemical cycles of elements and underpin marine food webs. Although their uncultured genetic diversity has been estimated by planetary-scale metagenome sequencing and subsequent reconstruction of metagenome-assembled genomes (MAGs), this approach has yet to be applied for eukaryote-enriched polar and non-polar phytoplankton communities. Here, we have assembled draft prokaryotic and eukaryotic MAGs from environmental DNA extracted from chlorophyll a maximum layers in the surface ocean across the Arctic Circle in the Atlantic. From 679 Gbp and estimated 50 million genes in total, we recovered 140 MAGs of medium to high quality. Although there was a strict demarcation between polar and non-polar MAGs, adjacent sampling stations in each environment on either side of the Arctic Circle had MAGs in common. Furthermore, phylogenetic placement revealed eukaryotic MAGs to be more diverse in the Arctic whereas prokaryotic MAGs were more diverse in the Atlantic south of the Arctic Circle. Approximately 60% of protein families were shared between polar and non-polar MAGs for both prokaryotes and eukaryotes. However, eukaryotic MAGs had more protein families unique to the Arctic whereas prokaryotic MAGs had more families unique to south of the Arctic circle. Thus, our study enabled us to place differences in functional plankton diversity in a genomic context to reveal that the evolution of these MAGs likely was driven by significant differences in the seascape on either side of an ecosystem boundary that separates polar from non-polar surface ocean waters in the North Atlantic.

## Introduction

The global ocean arguably harbours the largest microbial diversity on planet Earth. To reveal insights into global marine microbial diversity, which is also considered to be the biogeochemical engine of our planet, multiple large-scale international projects, of which TARA Oceans [1] might be the most significant, have been conducted over the past 10 years. The outcome of these projects has provided a step change in our understanding of marine microbial diversity especially in the surface ocean. One of the most important revelations from these initiatives was the realisation that we have significantly underestimated plankton diversity in the past because we were too reliant on culture-dependent methods [2]. As a consequence, some of the groups we thought of as being insignificant in the oceans turned out to be highly diverse with a major contribution to the global carbon cycle and marine food webs [3]. Furthermore, the significance of organism interactions and specifically symbiosis for cycling of energy and matter was revealed, with viral-host dynamics as the most impactful form of these biotic interactions [4, 5].

Linking functional microbial diversity, estimated by metagenomics and metatranscriptomics, with microbial activity as part of physico-chemical ecosystem properties shed light on how different microbial groups contribute to biogeochemical cycling of elements [1, 6, 7]. These results built the foundation for estimating how changing oceans due to global warming might impact the diversity and activity of ocean microbes [7, 8]. However, to fully explore the role of microbes and their interactions in changing environmental conditions, we must understand their metabolic capabilities in an evolutionary context [9, 10]. As the majority of marine microbes are unculturable and because genomic information is required to reconstruct their metabolic evolution, metagenome-assembled genomes (MAGs) offer a solution [11, 12]. Although most MAGs are not at the level of quality achieved through sequencing cultures of isolated strains, they provide genome-level insights into the microbial diversity of natural ecosystems. Due to their small size and structural simplicity, bacterial and archaeal genomes have preferentially been assembled from metagenomes [13, 14]. Hence, the majority of published MAGs are of prokaryotic nature and quite often eukaryotes are not even part of the underlying metagenomes due to selective filtration of microbial communities.

To the best of our knowledge, there are less than 20 reports on MAGs from oceanic habitats and all of them primarily report on prokaryotic genome reconstructions [14]. Nevertheless, these MAGs represent a new genomic resource and will help to analyse metagenome and metatranscriptome datasets as their analysis is largely limited by the availability of reference genomes. The latter particularly applies for eukaryotic microbes [12]. In addition to this phylogenetic bias, MAGs are also geographically biased because most of them have been reconstructed from tropical and temperate oceans [14]. However, a recent metagenomics study in the Arctic and Southern Oceans retrieved 214 prokaryotic MAGs [15], which appears to be the first study of this kind in polar sea water. Thus, the largest gap in our current knowledge on genomic diversity of uncultured oceanic microbes lies in polar oceans such as the Arctic and Southern Oceans and especially their microbial eukaryotes.

As polar marine ecosystems are under significant pressure because of global warming and as they disproportionally contribute to the global carbon cycle, there is an urgent need to reveal their genomic diversity [15]. Unlike in tropical and warm temperate oceans, primary production in polar oceans is mainly based on photosynthetic microbial eukaryotes such as diatoms, haptophytes, chlorophytes and prasinophytes [16–18]. Their genomes might be complex and can be large in size due to either genome duplications and/or the accumulation of repeats driven by the activity of transposable elements [19, 20]. Due to limited access to polar marine ecosystems and the temperature sensitivity of polar microbes, only very few genomes of these organisms have been sequenced so far [19–21], and comparative analyses of polar vs non-polar MAGs from uncultured prokaryotic microbes has only been reported once at least to the best of our knowledge [15]. To address this knowledge gap, we selected the Atlantic and adjacent Arctic Ocean for sequencing eleven surface ocean metagenomes from chlorophyll *a* maximum layers enriched for eukaryotic phytoplankton communities (size range 1.2 – 100 μm). A total of 679 Gbp representing 4.53 billion reads from 6 Arctic and 5 North Atlantic metagenomes resulted in the recovery of 140 MAGs including several draft genomes of microalgae. A comparative analysis of all MAGs revealed polar-specific metabolism and a demarcation between MAGs from Arctic vs temperate and subtropical North Atlantic surface waters. Thus, our study provides novel insights into uncultured genomic diversity of polar ocean microbes including differences to their non-polar counterparts.

## Material and Methods

### Sampling, DNA extraction and purification, sequencing and taxonomic identification

Samples were collected on two RV Polarstern (Alfred-Wegener Institute for Polar and Marine Research, Bremerhaven, Germany) expeditions described by [22] (Supplementary Data 1). Eleven samples were taken from chlorophyll a maximum layer of the surface ocean for metagenome sequencing. Six of these were stations within the Arctic circle, five in the temperate and subtropical North Atlantic. Arctic samples were collected on ARK-XXVII/1 (PS80) between 17^th^ June and 9^th^ July 2012; Atlantic samples were collected on ANT-XXIX/1 (PS81) between 1^st^ and 24^th^ November 2012. After water samples were pre-filtered with a 100 μm mesh to remove bigger zooplankton, they were filtered onto 1.2 μm Nucleopore membrane filters and stored at −80°C until further analysis. DNA was extracted using the EasyDNA Kit (Invitrogen, Carlsbad, CA, USA) with some adjustments. Cells were washed off the filter with pre-heated (65 °C) solution A from the kit and the supernatant was transferred into a new tube with one small spoon of glass beads (425-600 um, acid washed) (Sigma-Aldrich, USA). The samples were then vortexed three times in intervals of 3 seconds to break the cells. RNAse A was added to the samples and incubated for 30 min at 65 °C. The supernatant was transferred into a new tube and solution B from the kit was added followed by a chloroform phase separation and an ethanol precipitation. DNA was pelleted by centrifugation and washed several times with isopropanol, air dried and suspended in 100 uL TE buffer. DNA concentration was measured with a Nanodrop (Thermo Fisher Scientific, Waltman, MA, USA), samples snap frozen in liquid nitrogen and stored at −80C until sequencing. Description of the samples and associated metadata is available through the GOLD database [23].

All eleven samples were sequenced and assembled by the Joint Genome Institute (JGI), while annotation was performed using the Integrated Microbial Genomes & Microbiomes (IMG/M) pipeline [24, 25]. In summary, paired-end sequencing was performed on an Illumina HiSeq platform. BBDuk [26] was used to remove Illumina adapters, then BBDuk filtering and trimming applied. Reads mapping to the human HG19 genome with over 93% identity were discarded. Remaining reads were assembled with MEGAHIT [27]. The quality-controlled reads were mapped back to the assembly to generate coverage information using seal [28]. Some of these samples were later reassembled using SPAdes [29]. For eukaryotic binning, we used only the MEGAHIT assemblies. Prokaryote bins come from either the MEGAHIT or SPAdes assembly for that sample, though no sample had both assemblies binned.

Taxonomic classification and abundance estimation were performed using Bracken [30] and Kraken2 [31] (Supplementary Data 2). A custom Kraken2 database was constructed using all RefSeq genomes for bacteria, archaea, viruses, protozoa, fungi, as well as plants excluding embryophyta. Reads were taxonomically classified using Kraken, and abundance at the level of phylum was estimated with Bracken. Taxonomic classification had been performed as part of the IMG/M pipeline; three samples were processed in 2013/4 and the remainder in 2016/7. The taxa identified between the two groups showed clear differences, in the 2013/4 group a large amount of sequences assigned to the eukaryota and bacteria nodes rather than a more specific taxon. For this reason, we repeated taxonomic classification for all samples to ensure differences are not due to differences in reference databases or pipelines.

### Binning

The IMG/M pipeline identified a number of prokaryotic bins. Samples were binned by JGI as described in [24]. Briefly, each assembly was binned separately, using MetaBat [32] and a minimum contig size of 3000bp. Resulting bins were assessed for completion and contamination with CheckM [33] which also provides initial estimate of taxonomy. While eukaryotic sequences were not excluded from binning, all bins were labelled as archaea, bacteria or unknown by CheckM, prompting the distinct binning attempt for eukaryotes.

For eukaryotic binning, each assembly was binned separately, the process for binning one assembly is given below. Eukaryotic contigs were predicted with EukRep [12], which uses a linear support vector machine to classify sequences as eukaryotic or prokaryotic using k-mer frequencies. Coverage of the eukaryotic contigs was estimated by pseudoaligning the reads from each sample to the contigs using Kallisto [34]. Binning was performed using MetaBat [32] with the coverage information, and a minimum contig size of 1500bp. Completeness and contamination of resulting bins were assessed with BUSCO v3 [35], using the eukaryota_odb9 set of genes. Bins which were less than 50% complete were discarded from further analysis. Completion is defined as the percentage of expected single-copy genes from a selected gene set observed in a MAG, and contamination is defined as the percentage of single copy genes observed in two or more copies.

Names have been assigned to MAGs composed of the station they were binned from, a numerical identifier, and a suffix of either P to indicate they are from the IMG prokaryotic binning, or E to indicate they are from the eukaryotic binning. The numerical identifier is taken from the IMG portal; for eukaryotes the MAGs from a station are given ascending numbers starting from the MAG with highest completion.

Contigs in all MAGs, both prokaryotic and eukaryotic, were concatenated and reads pseudo-aligned back to this set of sequences representing all MAGs using Kallisto [32], to estimate the proportion of reads represented by the recovered MAGs.

### Phylogenetic placement

PhyloSift [36] was used to identify sequences homologous to the mostly-single copy genes in bins and reference genomes using the HMMs provided by PhyloSift. For eukaryotic reference genomes, all protists and green algae labelled representative from NCBI were used, as well as two diatom genomes (*Thalassiosira pseudonana*, *Phaeodactylum tricornutum*) taken from JGI. For prokaryotes, all genomes in the MarRef [37] database were included. Homologous sequences were located and the best hit retained when there were multiple. Viral marker genes were excluded. Marker genes present in less than 50% of the genomes (reference or MAGs) were not used in future steps of the analysis. Homologous sequences were aligned against the PhyloSift models, and alignments for all genes concatenated. FastTree [38] was used to build phylogenomic trees for the eukaryotic and prokaryotic alignments, using the general time reversible model option. The resulting trees were visualized with Interactive Tree of Life Viewer [39].

As additional evidence for taxonomy contigs from MAGs were searched against databases with BLAST [40] and each contig assigned a taxonomy using the MEGAN-LR algorithm [41]. Eukaryotes were searched against Marine Microbial Eukaryote Transcriptome Sequencing Project (MMETSP) [42], prokaryotes against NT. Selected groups of MAGs and reference genomes had ANI (Average Nucleotide Identity) calculated with pyani [43] using the BLAST-based ANIb method (Supplementary Data 7).

### Coverage

Coverage for each eukaryotic MAG was generated by aligning reads from each sample back to the bins using Bowtie2 [44] (Supplementary Data 3). Detection and mean coverage were calculated from these alignments using BedTools [45]. We considered a MAG not present in a sample if the detection was lower than 0.9.

### Functional annotation

Functional annotation for contigs was carried out as part of the IMG/M pipeline before binning. Protein coding genes were predicted using two prokaryotic gene prediction tools: Prodigal [46] and prokaryotic GeneMark.hmm [47]. For prokaryotes further gene prediction and annotation was not performed, the annotations for the contigs before binning were used. Gene Ontology (GO) terms for prokaryotic genes were generated using the mapping Pfam accession to GO terms maintained by InterPro. After binning, genes for contigs in eukaryotic MAGs were predicted ab initio using the eukaryote specific gene prediction tool GeneMark-ES [48] in self training mode with MAKER2 [49]. Predicted proteins were annotated using InterproScan 5 [50].

## Results

### Metagenome sequencing and annotation of contigs

Sampling stations have been named according to their geographical location in relation to the Arctic Circle. P-stations (polar) were located north and NP-stations (non-polar) south of the Arctic Circle in the North and South Atlantic (Figure 1a). In total, eleven stations were sampled (P1-6; NP1-5) and one metagenome was generated per station except for P3, which was used to sequence two metagenomes from two independent samples obtained from the chlorophyll *a* maximum layer. These two samples were labelled P3a and P3b. Sequencing all samples resulted in 4.53 billion reads totalling 679.25 Gbp, with each sample ranging between 46.79 Gbp and 67.37 Gbp. Assembling each station with MEGAHIT resulted in 42.10 million contigs totalling 23.02 Gbp.

**Figure 1.**
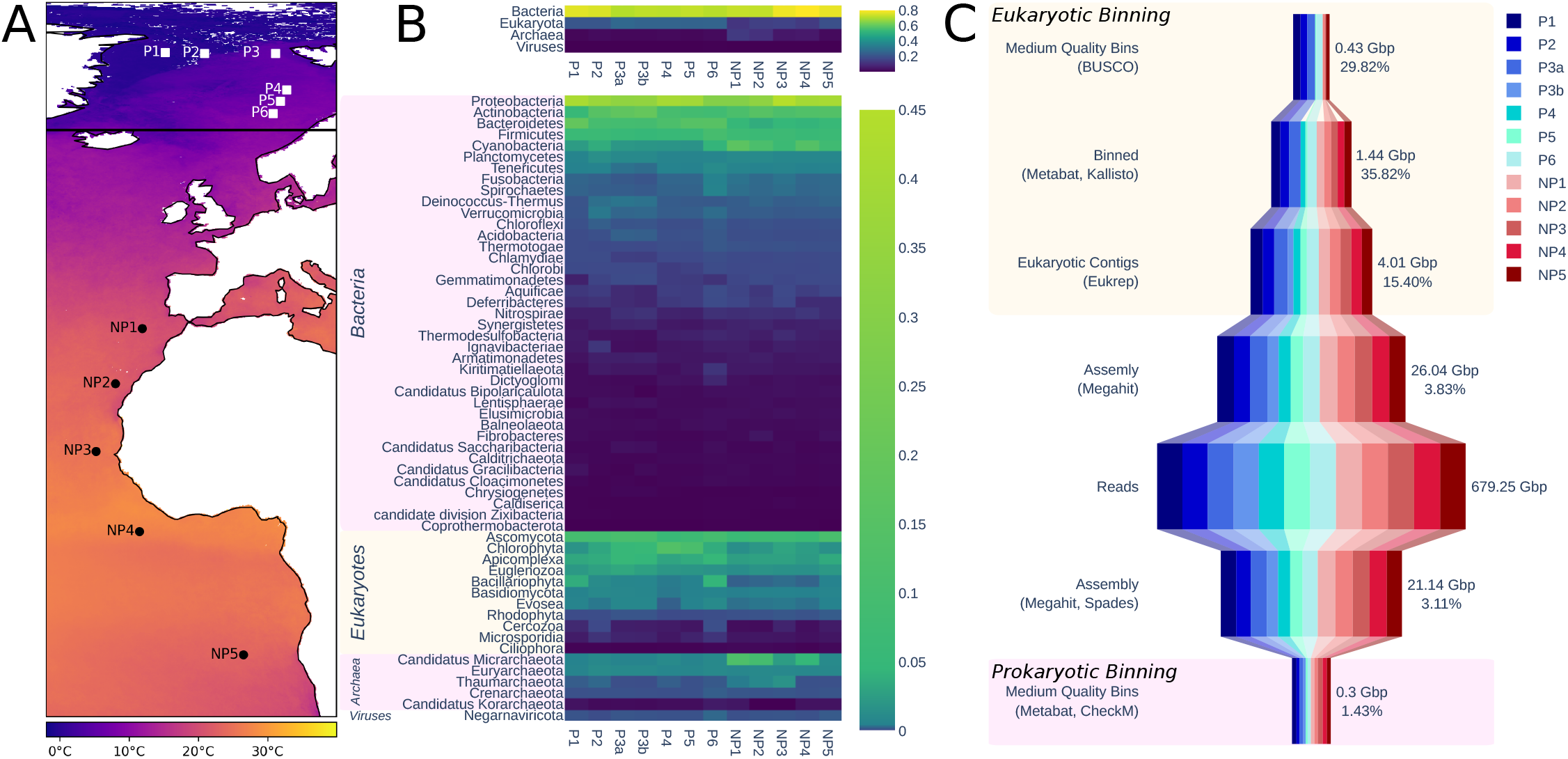
From left to right. A) Map showing sampling locations. Horizontal black line shows arctic circle. Colour indicates mean annual sea surface temperature for year of sampling [68]. B) Estimated relative taxonomic abundance. Top plot shows abundance summarised to the rank of superkingdom; bottom is summarised to rank of phylum. C) Summary of the size of data at different points of processing. Pink box indicates steps in the prokaryotic binning process, peach those in eukaryotic binning. Number of bases is the size of data in this step, percentage is the percentage of the data retained from the previous step.

Kraken2 taxonomically classified 365.85 million (15.74%) of the read pairs. Bracken abundance estimation at the level of superkingdom and phylum is shown in figure 1b. The most abundant read pairs were of bacterial origin followed by eukaryotes, archaea and viruses. On phylum level, read pairs from proteobacteria were most abundant with Ascomycota being the most abundant eukaryotic phylum followed by Chlorophyta. Generally, eukaryotes are more abundant in polar stations, contributing between 22% and 27% of the total abundance of reads, whereas they only contribute between 12% and 19% non-polar station. In non-polar stations with lower abundance of eukaryotes, there is a corresponding increase in the abundance of archaea. This is most pronounced in stations NP1 and NP2 (Figure 1b), where the most southern non-polar station NP5 appears to be more similar to polar stations. Photosynthetic microbes are present at all stations. However, photosynthetic eukaryotes such as chlorophytes and bacillariophytes generally have higher relative abundance in polar stations, whereas Cyanobacteria are more abundant in non-polar stations based on the relative contribution of reads.

IMG/M predicted 50.30 million genes in all sequenced metagenomes. Domains homologous to those in the Pfam database were found in 13.83 million (27.51%) of the predicted genes. Within samples, this proportion varied from 17.97% to 33%. The two samples from P3 had the lowest ratio of genes with homologous Pfam domains, both under 20%. Taxonomic affiliations were assigned to 17.74 million of the genes, of which 66% prokaryotic, 28% were eukaryotic, and 6% viral.

A majority (87%) of the identified Pfam domains were shared between Arctic and non-Arctic samples. However, the proportion of domains with unknown function was higher for domains uniquely found in either polar or non-polar stations than shared between them. Domains of unknown function constitute 16.55% of shared domains, but 23.76% and 29.71% in polar and non-polar metagenomes, respectively. Among domains unique to polar samples, 63.57% were identified in only one sample, and none were in all samples. For non-polar samples, only 43% of domains were present in only one sample, and 8.50% were in all samples.

### Metagenome-Assembled Genomes (MAGs)

#### 1) Binning and Quality

Metagenome binning generated 140 MAGs of medium and high quality, following the definitions for quality in [11]. Medium quality requires a completion of at least 50% and contamination less than 10%; high quality a completion of greater than 90% and contamination less than 5%, as well presence of certain rRNA genes and tRNAs. These MAGs represent 0.71 Gbp of assembled reads (Figure 1c), while 8% of all reads mapped back to the sequences contained in the combined 140 MAGs. Of all bins, 116 were classified as prokaryotes, 18 as eukaryotes and 6 were of unknown taxonomic affiliation according to default criteria applied by CheckM [33]. Among the prokaryotes, 111 were classified as bacteria, 5 as archaea, and CheckM was unable to classify six bins. These unknown bins had an average size of 2.90 Mbp and therefore likely represent prokaryotic genomes. Slightly more prokaryotic MAGs were retrieved from non-polar than polar metagenomes, 64 and 58, respectively. All prokaryotic MAGs from polar samples were classified to at least the phylum as either Bacteriodetes, Proteobacteria and Verrumicrobia. Verrucomicrobia were only recovered from polar metagenomes. Prokaryotic MAGs from non-polar metagenomes were more diverse and included the six unclassified MAGs with an average genome size of 2.90 Mbp. Classified MAGs included 19 assigned to the domain level of bacteria, 5 archaea, 6 Actinobacteria, and 3 Planctomycetes. MAGs of the latter 3 lineages were not recovered from any of the polar metagenomes.

Filtering the assembly for each sample to retain only eukaryotic contigs as predicted by EukRep resulted in 2,151,309 contigs totalling 4.01 Gbp. From these, we recovered 18 medium quality eukaryotic MAGs. Only four of these eukaryotic MAGs were retrieved from non-polar metagenomes. Taxonomy was assigned to the eukaryotic MAGs based on their placement in a phylogenomic tree; 7 placed with Mamiellophyceae reference genomes, 8 with Bacillariophyta, and the placement of the remaining 3 was less clear. All but one of the Bacillariophyta were recovered from polar metagenomes. Polar Mamiellophyceae MAGs placed in a clade with Micromonas, and the non-polar MAGs with Ostreococcus or Bathycoccus.

Prokaryotic MAGs have a mean completion of 74.30% and contamination of 2.68%. The MAG with highest completion is P1_21P at 99.62% and a contamination of 2.81%. Taxonomically this MAG was classified to the family level as Flavobacteriaceae. Prokaryotic MAGs have a median L50 of 11,402 bp and median size of 2.23 Mbp. Eukaroyotic MAGs have a mean completion of 59.23% and contamination of 1.28%, with a median size of 25.01 Mbp. Details of the MAGs are available in Supplementary Data 1. The MAG with the highest completion is P2_1E at 84.8%. All but one MAG is fragmented, with a median L50 of 5,229 bp. The exception is P2_1E, which contains many contigs longer than 50 kbp, the longest being 106 kbp.

Some phyla with relatively high abundance in our taxonomic classification based on reads had no MAGs retrieved. Ascomycota and Firimicutes have a high abundance, but no MAGs recovered, whereas MAGs were retrieved for the less abundant phyla Bacillariophyta and Verrucomicrobia. The evenness of abundance within phyla could lead to differing levels of coverage for genomes within phyla. A diverse phylum whose species are more evenly distributed would have low coverage compared to a less even phylum, affecting the ability to recover MAGs. To investigate the effect of intraphylum evenness on recovering MAGs, we calculated Simpson’s evenness measure using the number of reads assigned to species for all phyla of bacteria with a mean relative abundance equal to or higher than Verrucomicrobia, which was the least abundant phyla for which MAGs were recovered. Only bacteria were used, as for eukaryotic phyla other than Ascomycota there were few reference genomes available for the taxonomic classification database. Phyla from which MAGs were recovered had a lower mean evenness (0.31) than those from which no MAGs were recovered (0.50). A t-test showed this difference is significant for p = 0.01. Ascomycota similarly have a high mean evenness (0.74).

#### 2) Phylogenomic Placement

##### 2a) Prokaryotes

The phylogenomic tree for prokaryotes in figure 2b was constructed using concatenated alignments of 38 marker genes, a subset of those included in the PhyloSift package. Genomes of marine prokaryotes were retrieved from the MarRef database, for a total of 943 reference genomes (Supplementary Data 4) in addition to the 122 prokaryotic MAGs recovered in our study. The tree includes MAGs in which 50% or more of the selected marker genes were identified, a total of 88 of the MAGs. The largest group consists of 31 MAGs which placed within a clade with alpha-, beta-, and Gammaproteobacteria references. A further 24 placed with Bacteroidetes, of which 17 are in clades of Flavobacteriales.

**Figure 2a.**
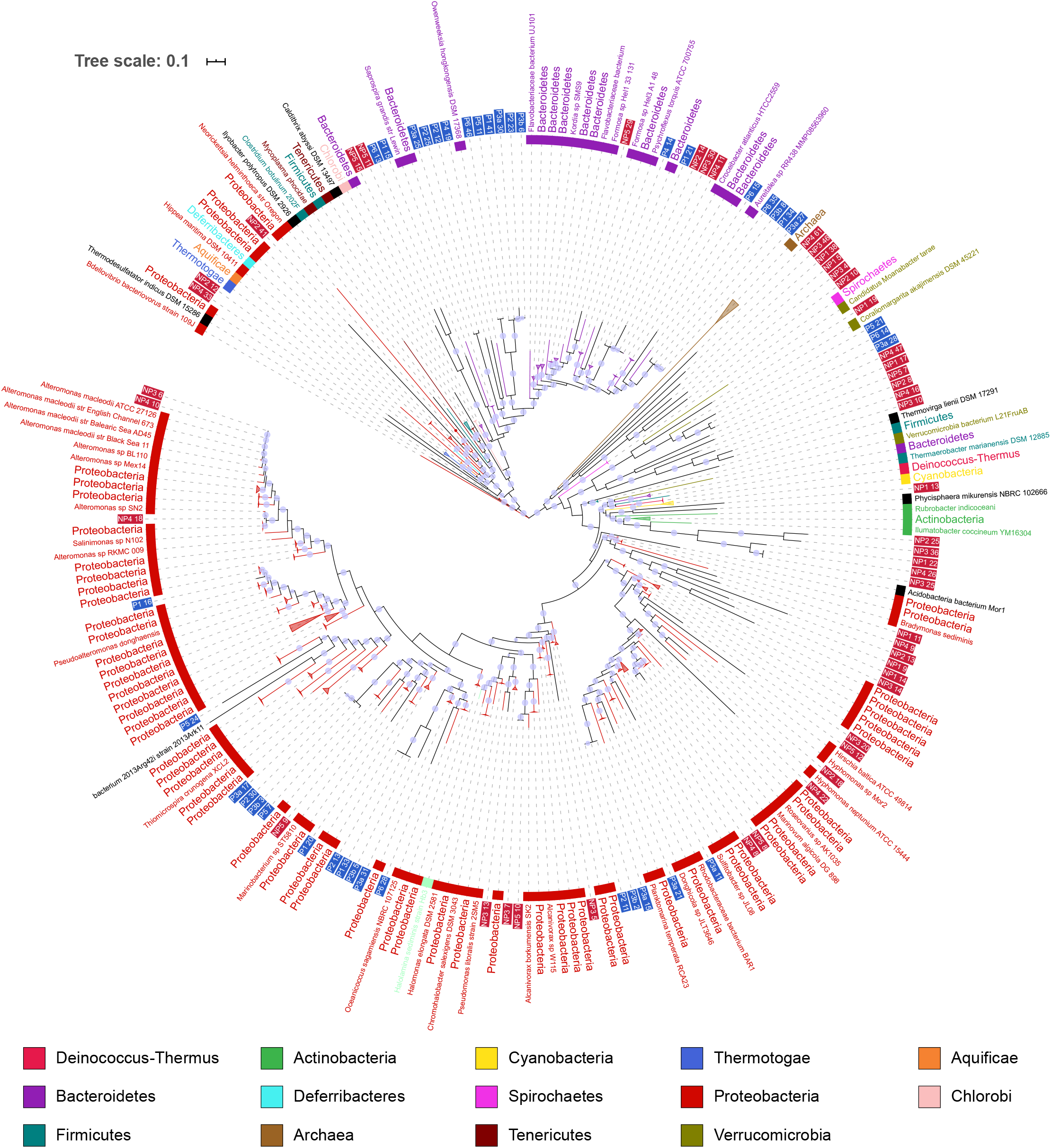
Phylogenomic tree including prokaryotic MAGs and MarRef reference genomes. Inner band color indicates taxonomy of reference genomes. MAG labels have blue background for polar MAGS, and a red background for non-polar. Clades which contained reference genomes all from the same taxonomic group in the legend have been collapsed. Local support values of greater than 0.75 are shown by a violet dot on branches.

**Figure 2b.**
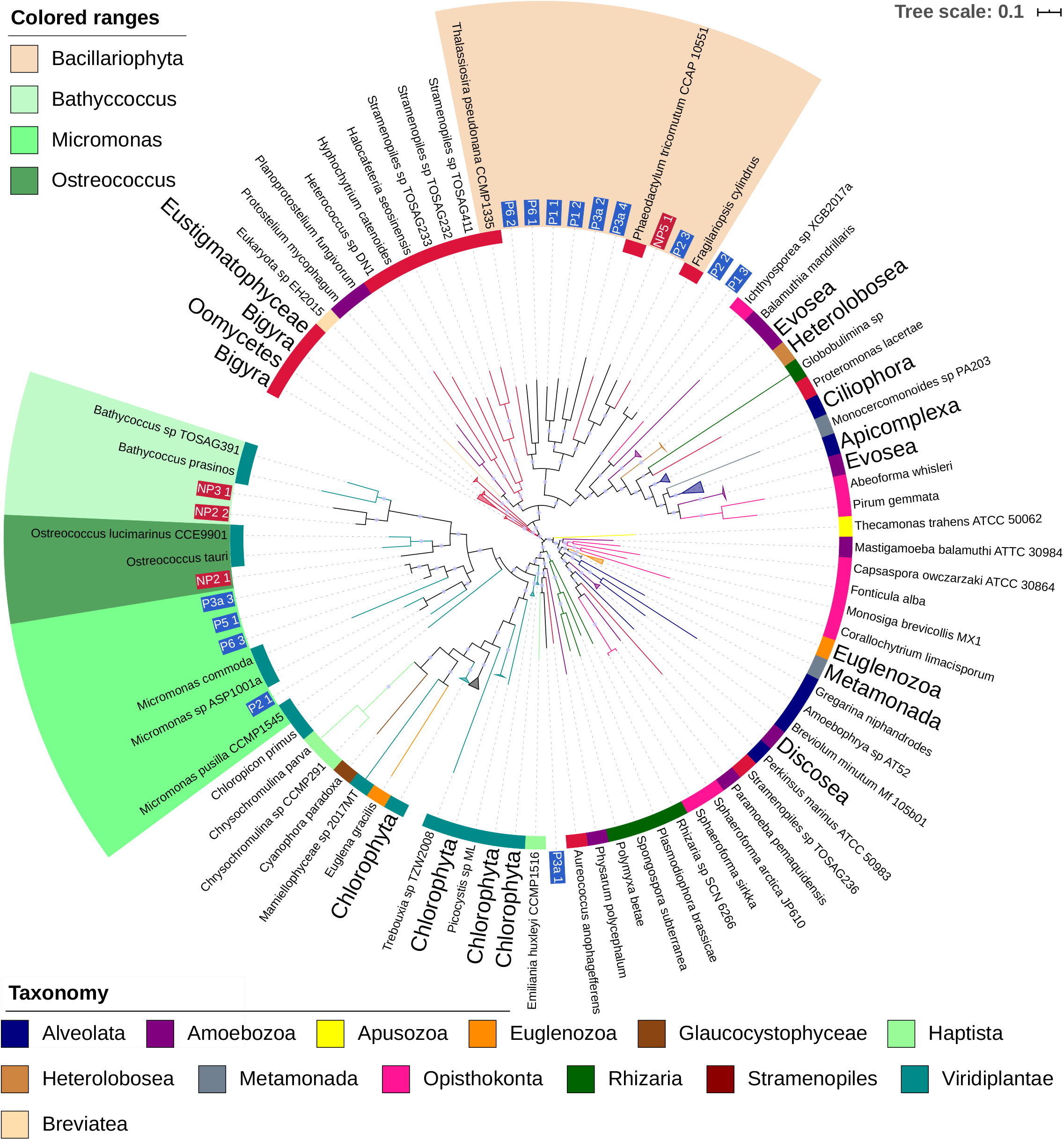
Phylogenomic tree including eukaryotic MAGs and reference genomes. Label and inner band color indicate taxonomy of reference genomes. MAG labels have blue background for polar MAGS, and a red background for non-polar. Clades which contained reference genomes all from the same taxonomic group in the legend have been collapsed. Colored ranges highlight clades where MAGs place with reference genomes of a consistent taxonomy. Local support values of greater than 0.75 are shown by a violet dot on branches.

The phylogenomic tree suggests taxonomy for some MAGS which were classified by CheckM either at the level of bacteria or had no classification. A group of 6 MAGs (NP1_11P to NP3_14P) form a clade close to Deltaproteobacteria references. Similarly, NP2_41P is placed among Epsilonproteobacteria and Oligoflexa. NP1_19P was classified as bacteria by CheckM, in the tree placed close to Puniceicoccales references and other MAGs which had been classified as Puniceicoccales by CheckM.

Some MAGs recovered from different stations appear closely related to one another. NP4_10P and NP3_6P are closely related to each other as well as to multiple *Alteromonas macleodii* strains. The reference genomes for *A. macleodii* can be split into those from surface and deep ocean [51], these MAGs have a greater than 95% ANI to three surface genomes, suggesting a species level relationship. The ANI between these MAGs and deep ocean *A. macleodii* is below 95%. This is supported by the assignment of contigs within the MAGs based on BLAST searches against the NT database, for both MAGs at least 89% of contigs are assigned to the *A. macleodii* node or a strain below it.

Other groups of MAGs display similarly close relationships to each other, but are more distant from reference genomes. Four polar MAGs which placed among Bacteroidetes, P6_35P, P3b_8P, P1_34P, and P3a_27P, share over 95% identity to each other, but less than that to their closest reference genome, an unclassified species of genus Aureitalea. The results of assigning contigs via BLAST searches is similarly mixed, most contigs being assigned to a mix of Flavobacterieaceae or uncultured bacterium. These four MAGs could represent members of the same novel species of Bacteroidetes.

There are few close relationships between polar and non-polar MAGs evident in the tree. The median distance from a polar MAG to the nearest polar MAG is lower than to the nearest non-polar MAG, and the same for non-polar to non-polar (Supplementary Data 5). In both cases the difference in medians is significantly different at p < 0.01 using Mood’s median test. One clade of Bacteroidetes is an exception, where polar MAG P1_21P appears closely related to NP2_14P, NP3_30P and NP4_11P. The closest reference is *Croecibacter atlanticus* which is in different clade. Pairwise ANI between these mags and the *C. atlanticus* reference genome is greater than 95%, suggesting these MAGs could represent genomes of the species *C. atlanticus*.

Some MAGs had been classified at a species level by CheckM where the phylogenomic tree does not suggest a similarly specific classification. MAGs P3a_28P, P6_14P, P5_21P, P2_21P, and P6_33P were classified as *Coraliomargarita akajimnesis* by CheckM. The first three placed closest to *C. akajimnesis* but with longer branches than observed between taxa from the same species elsewhere in the tree. The latter two lacked the amount of marker genes required to be included in the tree. Looking at the ANI also suggests these MAGs and *C. akajimensis* are not the same species, no pair shares above 95% ANI.

Three MAGs, NP1_5P, NP2_10P, and NP3_4P, were classified as planctomycetes by CheckM, but were more ambiguously placed in the phylogenomic tree. This may be a result of only one reference planctomycete genome, for *Phycisphaera mikurensi*, being used for tree construction.

##### 2b) Eukaryotes

The phylogenomic tree for eukaryotes in figure 2b was constructed using concatenated alignments of 57 marker genes, a subset of those included in the PhyloSift package. Representative genomes of microbial eukaryotes were retrieved from the National Centre for Biotechnology Information (NCBI) and JGI, for a total of 412 reference genomes (Supplementary Data 4) in addition to the 18 eukaryotic MAGs recovered in our study. Most MAGs placed in two clades, which contain all of the Bacillariophyta or Mamiellophyceae reference genomes. As branches within these clades are long, a more specific identification of these MAGs is difficult because of a lack of a sufficient number of reference genomes from eukaryotic marine microbes. Within the Mamiellophyceae clade, three MAGs (P6_3E, P5_1E, P3a_3E) are closely related to one another, but relationships to the reference genomes are more distant. Bacillariophyta-like MAGs appear to have more distant relationships (Figure 2). P2_2E and P1_3E are difficult to provide a taxonomy for. They placed close to each other, but distant from any reference genomes, and searches against MMETSP had no results for over 90% of contigs.

Mamiellophyceae-like MAGs appear to further divide into three clades containing reference genomes from the three genera Micromonas, Bathycoccus and Ostreococcus. Micromonas MAGs were only recovered from polar and Bathycoccus and Ostreococcus only from non-polar metagenomes. Some Mircomonas MAGs have high Average Nucleotide Identity (ANI) to each other or to reference genomes. For instance, MAG P2_1E has 99% ANI with Micromonas_1001a, a species reconstructed from an Antarctic metagenome [52]. Three MAGs appear highly similar: P6_3E, P5_1E and P3a_3E. ANI between these MAGs is 98% or higher, and 99% between P5_1 and P3a_3. However, this group do not share high ANI with any of the reference genomes used.

There is consistency in the taxonomic assignments of contigs within Mamiellophyceae MAGs at the phylum level. With the exception of NP2_1E, they have over 99% of their contigs assigned to Chlorophyta when searched against MMETSP as explained in Methods. The contigs that were not assigned to Chlorophyta were either assigned to the Eukaryota node, or had no BLAST hits. No contigs were assigned to other phyla. This suggests a consistent taxonomic origin for the sequences in these MAGs at least at the phylum level, rather than representing sequences which are not biologically related. Evidence from these BLAST searches supports the taxonomies suggested by the phylogenomic tree at the genus level; all Mamiellophyceae MAGs had at least 87% of their contigs assigned to the genus they placed with in the phylogenomic tree.

Within NP2_1E, there is less confirmatory evidence in the results of the BLAST searches, a greater number of contigs are not assigned a taxonomy or assigned to other phyla. This could represent either a MAG for an organism more distantly related to sequences available in the reference database, or increased contamination within the MAG. Contigs with no BLAST hits contributed 34.12% of all contigs. For those contigs that did have hits, 96% were assigned to Chlorophyta, which represents 63.44% of the total contigs in the MAG. Contigs assigned to other phyla constitute 2.13% of the total.

Eight MAGs placed in a clade with Bacillariophyta reference genomes, only 1 of which was non-polar. None of the MAGs appear close to the three reference genomes used. Some Bacillariophyta MAGs could be classified at genus level. For instance, MAG P2_3E had an ANI of 85.5% to *Fragilariopsis cynlindrus*, supporting their close placement. MMETSP contains sequences from Bacilliarophyta taxa which currently lack a complete genome, results from searching sequences in that MAGs against this database provided further evidence for taxonomy. Apart from MAG P3a_4E, all the MAGs in the Bacillariophyta clade had 85% or more of their assigned contigs classified at the level of phylum when searched against MMETSP as described in Methods. P3a_4E had a majority of contigs assigned to Bolidophyceae, a sister taxa to Bacillariophyta. As no complete genome from this class currently is available, this MAG may represent the first Bolidophyceae genome. An additional close placement was obtained for MAG P6_2E, for which ca. 84% of contigs were classified as *Leptocylindrus danicus.*

Many contigs in Bacillariophyta MAGs had no hits when searched against MMETSP with BLAST, a mean of 40.62% of contigs in Bacillarophyta MAGS had no hits. For comparison the mean percentage of contigs in Mamiellophyceae MAGs which had no hits in MMETSP was much lower, at 5.12%.

The MAG P3a_1E placed closest (ca. 73% ANI) to the Haptophyta *Emiliania huxleyi*. *E. huxleyi* is quite distant in the tree from the other two Haptophyta *Chrysocromulina parva* and *Chrysocromulina sp.* CCMP2291, which are from the Prymnesiales order. These two Prymnesiales placed as neighbouring leaves and showed 97% ANI. *E. huxleyi* and P3a_1 have much lower ANI with each other and the two Prymnesiales genomes. Searching contigs from P3a_1E against MMETSP, a majority of contigs were assigned to a range of Haptophyta taxa which included *E. huxleyi* among them, with most being assigned to *Phaeocystis antarctica*. Contigs were also assigned to several other phyla as well, possibly due to MAG contamination.

#### 3) Coverage of MAGs and associations

Next, we used read-coverage to analyse MAG distribution across polar and non-polar samples. Where less than 90% of bases had at least one read aligned to them, we considered a MAG to not be present at that station. The mean coverage of contigs in prokaryotic MAGs ranged between 2.73 and 375.07 with a mean coverage of 43.70. We used the mean coverage per million reads as an estimate of abundance of MAGs across stations (Figure 3). The binning process uses covarying coverage to group contigs into bins. Thus, for highly similar MAGs recovered from different assemblies, a similar pattern of coverage across sites would be expected. Four proteobacteria MAGs which appeared closely related in the phylogenomic tree, P3a_17P, P2_30P, P3b_3P, and P5_7P show this pattern strongly, with a very similar patterns of changing coverage from stations P1 to P6. Coverage of MAGs tends to form a gradient across stations with close geographic proximity. For the most part there is a clear demarcation between polar and non-polar MAGs. Of the 122 MAGs, 116 are only present in either polar or non-polar samples. MAGs detected in both tend to be detected in samples P1 and P2.

**Figure 3.**
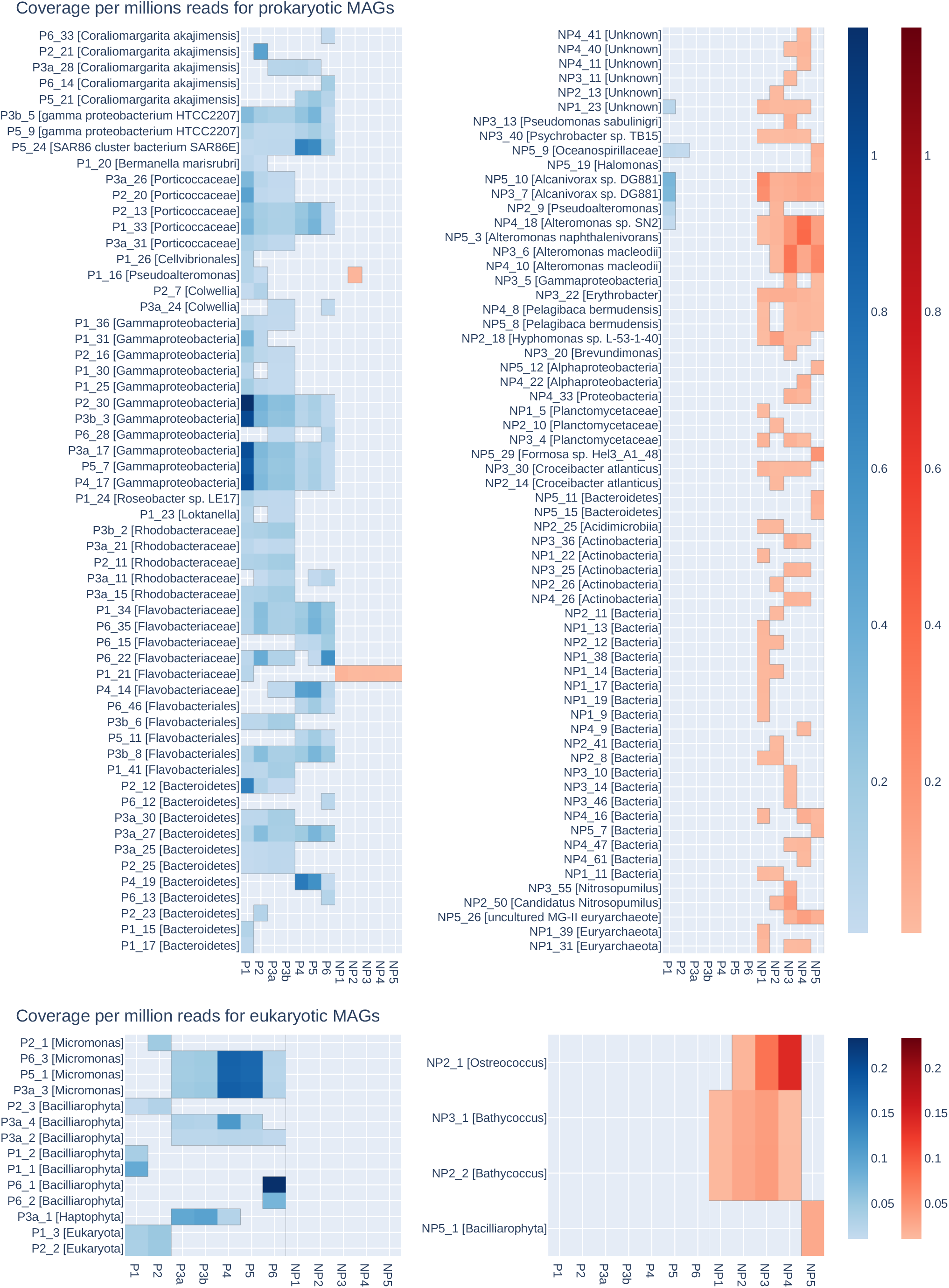
Mean coverage of each MAG in a given set of reads. Top shows prokaryotic MAGs, bottom eukaryotic MAGs. Coverage normalised to coverage per million reads. Coverage not shown where fewer than 90% of bases in a MAG had any read aligned. The left hand heatmaps show MAGs recovered from polar assemblies, right hand shows those recovered from non-polar assemblies. Coverage in reads from polar stations is shown in a blue scale, coverage in non-polar stations shown in a red scale. MAGs are ordered by taxonomy. Each MAG has been given a taxonomic label of the most specific rank to which taxonomy had been determined.

For eukaryotic MAGs, mean coverage ranged between 4.35 and 87.24, with a mean coverage lower than that of prokaryotes at 22.60 (Figure 3). Again, highly similar Micromonas MAGs P6_3E, P5_1E and P3a show very similar patterns coverage from stations P3 to P6. Coverage of MAGs tends to form a gradient across stations with close geographic proximity. There is clear demarcation between polar and non-polar eukaryotic MAGs, as no MAG was found to be on both sides of the Arctic Circle.

Most of the Bacillariophyta MAGs were present at only one or two stations maximum whereas Mamiellophyceae MAGs were more widespread such as P3a_2E and P3a_4E. The one non-polar Bacilliarophyta MAG is present only in station NP5, the southernmost of the non-polar stations. The closely related MAGs P2_2E and P1_3E which had not been assigned a taxonomy are present only in the two stations they were recovered from. Potential Haptophyte P3a_1E is present in two polar stations, and most abundant at P3, where the Mamiellophyceae MAGs are less abundant.

In both prokaryotes and eukaryotes, the MAGs which could not be assigned a taxonomy, assigned either as Unknown, Bacteria or Eukaryota, are mostly observed in 3 or fewer stations, with low coverage. The only exception is Prokaryote NP1_23P is an exception, detected in 5 stations.

Some associations were observed between the coverage of pairs of eukaryotic and prokaryotic MAGs (Supplementary Data 6). Ostreococcus MAG NP2_1E showed a roughly linear correlation to four prokaryotic MAGs (NP5_3P, NP 4_18P, NP4_47P, and NP4_40P); the first two were classified as Alteromonas species, the third only to the level of Bacteria, the final as a prokaryote. The three Micromonas MAGs (P3a_3E, P5_1E, P6_3E) show some association to P5_24P (SAR86 cluster bacterium) and P4_14P (Flavobacteriaceae), with a group of stations where both are observed at similarly low levels of coverage, and another group where both are present at higher.

#### 4) Functional annotation of MAGs

A PCA analysis of the Pfam abundance in each MAG (Figure 4) largely shows separation into taxonomic groups, supporting the broad classifications drawn from the phylogenomic tree. Clustering by taxonomy is stronger for eukaryotes than prokaryotes. The two large groups of Bacillariophyta and Mamiellophyceae are clearly separated, with the possible Bolidophyceae P3a_4E closer to P3a_1E the potential Haptophyte. Some prokaryotic groups form clear clusters, such as Bacteriodetes and Actinobacteria, while others are more spread such as the Proteobacteria. Most of the MAGs without an assigned taxonomy cluster to the right of the plot. The number of Pfams observed in these groups is shown in figure 3, for the whole population before binning, and for eukaryotic and prokaryotic MAGs. The whole population showed a majority of Pfams present in all studied geographical regions, suggesting a widely distributed shared core of functions. Among functions unique to either side of the Arctic Circle, prokaryotic MAGs had many more unique functions in non-polar waters whereas eukaryotes had more unique functions at polar waters. Only 4 eukaryotic MAGs had been recovered from non-polar metagenomes. This imbalance could partially explain the high number of functions unique to polar eukaryotic MAGs. Prokaryotic MAGs were more balanced across the Arctic Circle, 65 from non-polar and 58 from polar stations.

**Figure 4.**
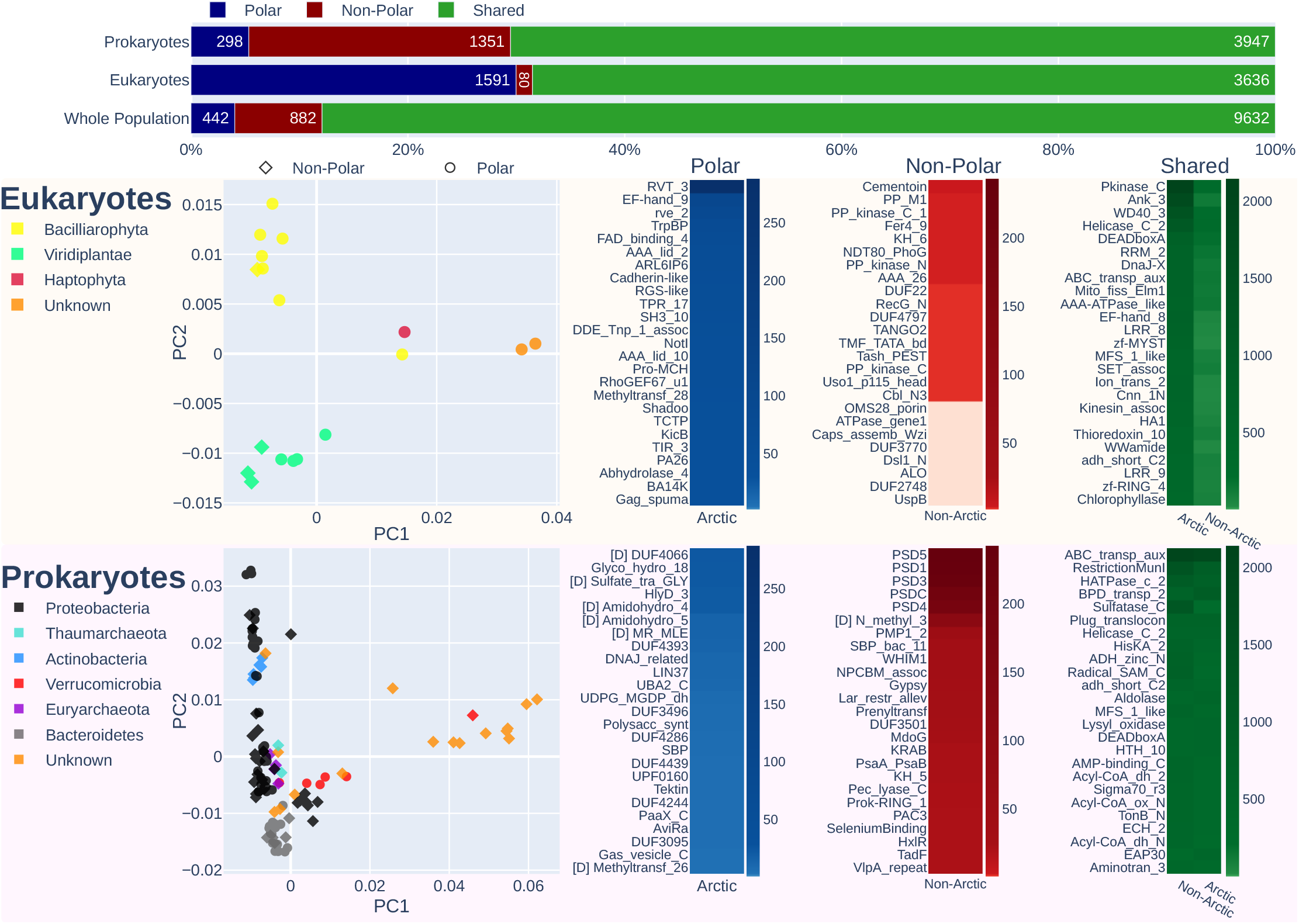
In the top, each horizontal bar shows how many Pfam accessions are found only in polar sequences (blue), only in non-polar (red) and found in both (green). This is shown for prokaryotic MAGs, eukaryotic MAGs, and for the whole population metagenome before binning. Below, the peach box shows information for eukaryotic MAGs, and the pink box for prokaryotic MAGs. For each, leftmost is a PCA plot of the proportion of Pfam in each MAG, with colours showing the taxonomy of each point. To the right heatmaps indicate the most abundant Pfams unique to polar (blue), non-polar (red) or shared (green). Pfams which are now dead families or merged since annotation are indicated with a D by the name.

Four of the five most abundant Pfam families unique to non-polar prokaryotic MAGs are PSD1, 3, 4, 5 & C. These are domains of unknown function shared by cytochrome-like proteins in the planctomycete species *Rhodopirellula baltica*. Three MAGs classified as planctomycetes were recovered, all from non-polar metagenomes. These domains were found in 24 of the 65 non-polar prokaryotic MAGs, which were assigned to a wide taxonomic range: Acidimicrobiia, Actinobacteria, Alphaproteobacteria, Gammaproteobacteria, Planctomycetaceae, and MAGs, which were classified as either bacteria or unclassified. All five proteins are typically found together in MAGs, only NP2_9P contained one (PSD3) without the others being present. The three Planctomycete MAGs are richer in these domains than others, accounting for 27.59% of the non-polar unique PSD domains. Along with the 15 MAGs classified only at the level of bacteria making up 67.93% these two groups account for a majority of our PSD encoding MAGs. The related domains PSCyt2, PSCyt3, PSD2 were shared between polar and non-polar MAGs, being found in four polar MAGs all identified as Puniceicoccaceae.

Eukaryotic MAGs have more Pfams unique to polar environments. The most abundant domain only found in polar MAGs is RVT_3, a domain believed to part of a retrotransposon found in plants [53]. RVT_3 was most abundant in two of the Bacillariophyta MAGs P6_1 with 125 and P6_2 with 144. This domain has been observed in complete genomes for Bacillariophyta, but in lower numbers. Another transposase related domain, rve_2, was observed in a high numbers of genes in P6_1E and P6_2E. Rve_2 is an integrase catalytic domain, present in transposase proteins as well as catalysing reactions involved in the integration of viral genomes into host genomes.

## Discussion

### Binning and retrieving of MAGs from phytoplankton metagenomes

Despite enrichment for larger eukaryotic phytoplankton, the most dominant group of organisms in the sequenced metagenomes was the group of bacteria (Figure 1). This might not only reflect their overall dominance but also their close association and importance for the growth of many eukaryotic phytoplankton species as previously shown in field and laboratory experiments [54, 55]. However, reads from eukaryotes represent the second most abundant group of organisms followed by archaea and viruses. As previously revealed by TARA Oceans metagenomes, the most prevalent groups of bacteria in the surface ocean are Proteobacteria, Actinobacteria and Bacteroidetes [1, 14]. We did not find any significant differences in their read abundance between polar and non-polar metagenomes, which confirms their ubiquity. Surprisingly, Ascomycota was the most abundant group of microbial eukaryotes in our metagenomes without significant geographic differences, which suggests that these fungi are very ubiquitous [56, 57]. However, this could be partially due to the greater number of Ascomycota reference sequences available; in the database used for taxonomic classification over half the eukaryotic minimisers mapped to Ascomycota species. They are known to be commensals or parasites of many different pelagic species including algae and animals but their roles and ecological functions in the surface ocean are far from understood [58]. Previous surveys based on phylogenetic marker genes revealed their diversity in the surface ocean including the Arctic [59], but our dataset suggests that at least in samples enriched for microbial eukaryotes, they appear to be more prominent than any autotrophic prokaryotes and eukaryotes regardless of geographic location. For reads from photosynthetic microbes, there appear to be geographical preferences relative to either side of the Arctic circle. Reads from cyanobacteria were more abundant in non-polar waters whereas reads from chlorophytes and bacillariophytes were more abundant north of the Arctic circle. All other groups identified in our metagenomes including the groups of Apicomplexa and archaea had a more patchy geographical distribution.

The retrieving of MAGs from metagenomes was not always in correspondence with the abundance of reads from specific taxonomic groups. This mismatch is potentially caused by a combination of factors. Sequencing depth, read length and the quality of reads most likely play a significant role in relation to genome size and complexity. The latter two factors might be the reason why we did not retrieve any MAGs from Apicomplexa such as dinoflagellates. Intraphylum diversity most likely plays a role, too [60]. For instance, it is known that the phylum Ascomycota is very species rich encompassing relatively small genomes [57]. Although Ascomycota appear to be the most abundant eukaryotic phylum in our dataset, we were not able to assemble any MAGs from this phylum because the read coverage for individual MAGs to be retrieved might have been insufficient. Populations with low diversity and high coverage have been observed to improve the quality of MAGs recovered by Metabat [61]. Thus, our results suggest that intraphylum evenness may affect the recovery of MAGs. Viridiplantae on the other hand are less diverse and especially members from the Prasinophytes have small genomes and are abundant in the surface ocean [62], which might explain why we retrieved several MAGs from different classes. Overall, completion of these MAGs is lower than the eukaryotic MAGs reported in prior eukaryotic binning studies [12, 63].

The proportion of prokaryotic MAGs recovered from different phyla are similar to those found in a larger study of oceanic diversity which recovered 2,631 [14]. In both the largest number of MAGs was from proteobacteria, followed by Bacteroidetes. Our results did not recover MAGs from some phyla which were recovered in high numbers by [14]. For example, 167 Chloroflexi MAGs were recovered in [14], where none of the MAGs we recovered were identified as Chloroflexi. Despite appearing to be one of more abundant phyla in our sample, neither binning effort identified Firmicutes MAGs, although similar studies using human gut data have [13, 14].

### MAG distribution, diversity, and abundance

The very pronounced demarcation between polar Arctic and non-polar Atlantic MAGs (Figure 3) for both prokaryotes and eukaryotes likely is a consequence of how major differences between both climatic regions have shaped the evolution and diversity of phytoplankton communities [8, 64]. The most significant difference is the seasonal presence of sea ice north of the Arctic circle. Freezing and melting of the surface ocean has a major impact on thermohaline mixing and therefore a variety of key environmental factors (e.g. light, nutrients) in addition to the overall low temperature in polar waters shaping the evolution, diversity, and activity of pelagic organisms [8, 64]. It has previously been proposed that the seascape boundary between seasonally mixed and permanently stratified waters at around the 15°C annual-mean isotherm separates global differences in oceanic primary production [64]. This isotherm also appears to be responsible for the latitudinal partitioning of microbiome compositions based on global ocean metatranscriptomes and metagenomes [8]. As this isotherm is separating our polar and non-polar communities although the polar sampling stations were further north of the 15°C annual-mean isotherm, it is likely causative for the strong demarcation between polar and non-polar MAGs. This suggests that this ecological boundary does not only affect the distribution of individual sequences in complex meta-omics datasets but also the diversity and evolution of genomes. However, some prokaryotic MAGs (e.g. P1_16P, P1_21P, NP_23P, NP_10P) have crossed this boundary, which might indicate the presence of locally adapted ecotypes. None of the eukaryotic MAGs has been found on both sides of the boundary, which suggests that the environmental differences might have had a stronger impact on diversification and therefore adaptation and evolution. These MAG-specific geographical distribution patterns are reflected in cross-kingdom co-occurrences between eukaryotes and prokaryotes in these phytoplankton communities (Supplementary Data 6). The co-occurrence patterns we identified were limited to either the Arctic or Atlantic side of the ecological boundary. Thus, none of them was crossing it, which indicates that co-evolution under significantly different environmental conditions was likely driving the formation of these associations. This, to the best of our knowledge, is the first example of how ecological boundaries in the seascape not only influence the spatial heterogeneity of sequences but genomes from co-occurring species in complex phytoplankton communities.

### Polar vs non-polar metabolism in MAGs

The separation of Pfams into taxonomic groups confirms the overall taxonomic placements of the MAGs based on concatenated phylogenetic approaches, even though the Pfam separation is less clear for prokaryotes (Figure 4). The latter might be caused by a higher proportion of genetic exchange between bacterial strains compared to their eukaryotic counterparts. Although whole population metagenomics already provided some evidence that there are region-specific Pfams (Figure 4), only the specific analysis of MAGs has revealed significant differences between prokaryotes and eukaryotes in terms of their genetic repertoire in relation to either side of the Arctic circle. The reason for eukaryotic MAGs to have more unique Pfams in polar waters and vice versa for prokaryotes remains enigmatic but suggests that identical environmental conditions and therefore similar selection pressures would impose differences in how prokaryotic and eukaryotic genomes evolve in the surface ocean. It appears that for eukaryotes, a dynamic surface ocean with seasonal mixing and sea-ice formation requires genomes to diversify because of the high abundance of transposable elements [19]. In contrast, prokaryotic MAGs in the same environment were characterised by a high abundance of domains of unknown function. Non-polar environments characterised by higher temperatures, stratified waters and weaker seasonality appear to enrich for PSD domains that are shared by chytochrome c – like proteins for electron transport as part of the respiratory chain in prokaryotes (Figure 4). This potentially suggests that respiratory activity is enhanced in non-polar prokaryotes compared to their polar counterparts, which would be expected according to the positive relationship between temperature and metabolic activity [65]. Interestingly, Pfams related to phosphate acquisition and metabolism in addition to Pfams involved in iron metabolism and electron transport were among the most enriched domains in non-polar eukaryotic MAGs. The relatively low nutrient concentrations in these stratified waters might only allow eukaryotes to thrive if they have developed mechanisms for the efficient uptake of nutrients [66, 67]. Smaller-sized prokaryotes with streamlined genomes usually outcompete eukaryotes in these environments as their nutrient demand is lower [66].

### Conclusions

Our study has revealed that surface ocean ecosystem boundaries separating significantly different oceanic provinces impact the evolution of prokaryotic and eukaryotic genomes in complex communities. They also appear to shape the nature of cross-kingdom co-occurrence patterns. Thus, MAG-based analyses of phytoplankton communities not only offer the identification of novel genomic resources, they might reveal unifying concepts responsible for how differences in ecosystem properties shape the genomes of their inhabitants and even species associations, which underpin the evolution of complex microbial communities.

## Acknowledgements

This work was supported by the Natural Environmental Research Council [NE/N012070/1]. The work conducted by the U.S. Department of Energy Joint Genome Institute is supported by the Office of Science of the U.S. Department of Energy under contract no. DE-AC02-05CH11231. The authors would like to thank the following collaborators from the Joint Genome Institute: A. Clum, A. Copeland, B. Foster, Br. Foster, M. Huntemann, N. N. Ivanova, N. C. Kyrpides, E. Lindquist, S. Mukherjee, K. Palaniappan, and T.B.K. Reddy. Sea surface temperature data in figure 1 taken from NASA Goddard Space Flight Center, Ocean Ecology Laboratory, Ocean Biology Processing Group; (2014): Moderate-resolution Imaging Spectroradiometer (MODIS) Aqua 11μm Day/Night Sea Surface Temperature Data; 2014 Reprocessing, NASA OB.DAAC. doi: data/10.5067/AQUA/MODIS/L3B/SST/2014. Accessed on 09/01/2020. A. Duncan was supported by a PhD studentship from the NEXUSS Centre for Doctoral Training (Environmental Research Council and the Engineering & Physical Sciences Research Council, UK)

## Supplementary Information

All MAGs, tables of predicted functions, and trees summarising taxonomic placement of contigs from BLAST searches have been made openly available on figshare at http://doi.org/10.6084/m9.figshare.c.5017517. Prokaryotic MAGs are additionally available via the IMG website, using the IMG Bin IDs provided in Supplementary Data 1.

List of files in Supplementary Information:

1. summary_statistics.xlsx: Summary details of data. Includes worksheets giving completion and contamination of MAGs.
2. taxonomy.tar.gz: Number of reads assigned to each taxon by Bracken at phylum and class level for all stations, in tab-separated format. Kraken 2 output provided for all stations in tab-separated format. Calculations for intraphylum evenness are included.
3. coverage.csv: Mean coverage and detection of MAGs for each set of reads. This data is not normalised for number of reads in each set of reads.
4. trees.tar.gz: For eukaryotes and prokaryotes, phylogenomic tree in Newick format, list of the Phylosift marker genes included when building tree, and details of reference genomes included in tab-separated format.
5. Tree distances between MAGs and the closest Polar and Non-Polar MAGs, displayed as box plots. Statistics between pairs are p-values from Mood’s median test for difference in sample medians.
6. associations.pdf: Scatter plots showing normalised coverage at stations between eukaryotic and prokaryotic MAGs where some association was observed.
7. ani.tar.gz: Average Nucleotide Identity plots and data in tab-separated format for related groups of MAGs and reference genomes.

