## Supplementary material for "Metagenome-assembled genomes of phytoplankton communities across the Arctic Circle": Suppl. Data 2: phylum_evenness.html


In [3]:

```
import pandas as pd
import glob
import os
```

In [4]:

```
# Function to load data for a specific Phylum
FILE_LOC = "station_data"
def load_phylum(phylum):
    data = None
    for file in glob.glob(f'{FILE_LOC}/*.kreport2'):
        if "bracken" in file:
            continue
        if not os.path.isfile(file):
            continue
        with open(file) as f:
            phy_lines = []
            found = False
            for line in f:
                parts = [x.strip() for x in line.strip().split('\t')]
                # Check if this is the start of the phylum
                if found and parts[3] == 'P':
                    found = False
                    break
                if parts[3] == 'P' and phylum.upper() in parts[5].upper():
                    found = True
                    continue
                if found and parts[3] == 'S':
                    parts[1] = int(parts[1])
                    phy_lines.append(parts)
        # To dataframe
        df = pd.DataFrame(phy_lines, columns=['pc', 'summed', 'direct', 'rank', 'taxid', 'name'])
        df['station'] = [os.path.basename(file).split('.')[0]] * len(df)
        if data is None:
            data = df
        else:
            data = data.append(df)
    piv = data.pivot(index='name', columns='station', values='summed')
    return piv

# To calculate simpsons e for evenness
import skbio.diversity.alpha as alpha
def evenness(x):
    return alpha.simpson_e(x[x>0])
def evals(datba):
    return data.fillna(0).apply(evenness)
```

In [8]:

```
# Get E for all top phyla
PHYLA = {
    'mag' : ['Proteobacteria', 'Actinobacteria', 'Bacteroidetes', 'Verrucomicrobia'],
    'no_mag' : ['Firmicutes', 'Cyanobacteria', 'Planctomycetes', 'Tenericutes', 
                'Fusobacteria', 'Spirochaetes', 'Deinococcus-Thermus']
}

evens = []
for name, phyla in PHYLA.items():
    elist = []
    for phylum in phyla:
        data = load_phylum(phylum)
        e = evals(data)
        elist += list(e.values)
        print(phylum, e)
        print(f'{phylum} no taxa: {len(data)}')
    evens.append((name, elist))

import statistics
print('')
print(f'MAG    | mean: {statistics.mean(evens[0][1])}, variances: {statistics.variance(evens[0][1])}')
print(f'No MAG | mean: {statistics.mean(evens[1][1])}, variances: {statistics.variance(evens[1][1])}')

from scipy import stats
stats.ttest_ind(evens[0][1], evens[1][1], equal_var=True)
```

```
Proteobacteria station
NP1    0.057917
NP2    0.052425
NP3    0.009802
NP4    0.013371
NP5    0.012972
P1     0.016511
P2     0.010243
P3a    0.036875
P3b    0.040555
P4     0.004326
P5     0.004333
P6     0.048157
dtype: float64
Proteobacteria no taxa: 2431
Actinobacteria station
NP1    0.502894
NP2    0.521922
NP3    0.515067
NP4    0.531255
NP5    0.515261
P1     0.438075
P2     0.416139
P3a    0.072399
P3b    0.103223
P4     0.522706
P5     0.521113
P6     0.550547
dtype: float64
Actinobacteria no taxa: 867
Bacteroidetes station
NP1    0.149906
NP2    0.318397
NP3    0.331226
NP4    0.405500
NP5    0.496813
P1     0.080186
P2     0.198658
P3a    0.098436
P3b    0.097698
P4     0.215568
P5     0.202539
P6     0.311343
dtype: float64
Bacteroidetes no taxa: 409
Verrucomicrobia station
NP1    0.737572
NP2    0.687550
NP3    0.698848
NP4    0.713466
NP5    0.689040
P1     0.552017
P2     0.308046
P3a    0.503293
P3b    0.468655
P4     0.512278
P5     0.495593
P6     0.328361
dtype: float64
Verrucomicrobia no taxa: 17
Firmicutes station
NP1    0.297653
NP2    0.279934
NP3    0.290895
NP4    0.314048
NP5    0.239449
P1     0.307838
P2     0.294339
P3a    0.247276
P3b    0.267527
P4     0.322330
P5     0.293195
P6     0.307740
dtype: float64
Firmicutes no taxa: 862
Cyanobacteria station
NP1    0.011522
NP2    0.013340
NP3    0.021670
NP4    0.014133
NP5    0.013953
P1     0.451603
P2     0.302178
P3a    0.440570
P3b    0.432253
P4     0.375679
P5     0.419130
P6     0.381288
dtype: float64
Cyanobacteria no taxa: 141
Planctomycetes station
NP1    0.614328
NP2    0.478144
NP3    0.513168
NP4    0.418917
NP5    0.321721
P1     0.236382
P2     0.042976
P3a    0.282605
P3b    0.310176
P4     0.293775
P5     0.223481
P6     0.590595
dtype: float64
Planctomycetes no taxa: 85
Tenericutes station
NP1    0.805189
NP2    0.815160
NP3    0.823908
NP4    0.815784
NP5    0.816170
P1     0.747758
P2     0.292595
P3a    0.699830
P3b    0.666544
P4     0.794692
P5     0.778046
P6     0.291311
dtype: float64
Tenericutes no taxa: 122
Fusobacteria station
NP1    0.550864
NP2    0.536003
NP3    0.546446
NP4    0.541688
NP5    0.550477
P1     0.538861
P2     0.374171
P3a    0.574732
P3b    0.578464
P4     0.570725
P5     0.552386
P6     0.486583
dtype: float64
Fusobacteria no taxa: 24
Spirochaetes station
NP1    0.595246
NP2    0.606918
NP3    0.583280
NP4    0.596251
NP5    0.634717
P1     0.601311
P2     0.648659
P3a    0.697585
P3b    0.709289
P4     0.608994
P5     0.624814
P6     0.493513
dtype: float64
Spirochaetes no taxa: 54
Deinococcus-Thermus station
NP1    0.818081
NP2    0.802373
NP3    0.805743
NP4    0.792395
NP5    0.808035
P1     0.614654
P2     0.680512
P3a    0.464230
P3b    0.759570
P4     0.816835
P5     0.810776
P6     0.718760
dtype: float64
Deinococcus-Thermus no taxa: 31

MAG    | mean: 0.3149807820784989, variances: 0.05721219847277323
No MAG | mean: 0.49650876462267446, variances: 0.05172962842549736
```

Out[8]:

```
Ttest_indResult(statistic=-4.328945027327323, pvalue=2.9654413187889486e-05)
```

In [5]:

```
# Evenness for Ascomycota
asco = load_phylum('Ascomycota')
asco_e = evals(asco)
print('Ascomycota', e)
print(f'Mean e: {e.mean()}')
```

```
Ascomycota station
ArcOceMetagenome_2_FD    0.680512
ArcOceMetagenome_3_FD    0.464230
ArcOceMetagenome_FD      0.614654
AtlANTMetagenome_2_FD    0.802373
AtlANTMetagenome_FD      0.818081
FraStrMetagenome_FD      0.718760
NorSeaMetagenome_2_FD    0.810776
NorSeaMetagenome_FD      0.816835
SouAtlMetagenome_2_FD    0.808035
SouAtlMetagenome_FD      0.759570
TroAtlMetagenome_2_FD    0.805743
TroAtlMetagenome_FD      0.792395
dtype: float64
Mean e: 0.7409968844214251
```

In [ ]:
