## Supplementary material for "Metagenome-assembled genomes of phytoplankton communities across the Arctic Circle": Suppl. Data 2: prokaryote_coverage.html


In [1]:

```
import pandas as pd
import matplotlib.pyplot as plt
import papername
import os
import seaborn as sns
import numpy as np
```

In [2]:

```
# Define methods for reading and converting data
# Method to read coverage information
def read_abundance(path, stations):
    # Returns either a DataFrame or an exception
    data = pd.read_csv(path)
    # Trim any unwanted stuff from the station column
    data['portal'] = data['portal'].apply(lambda x: os.path.basename(x).split('.')[0])
    # Merge with station details if they exist
    if not stations is None:
        data = pd.merge(data, stations, on='portal', how='left')
    # apply some sorting
    data.sort_values(by=['mag', 'station'])
    return data

def read_stations(path):
    #Returns a table of station details
    data = pd.read_csv(path)
    return data.set_index(data.columns[0])

# Methods for comparing coverage
def coverage_matrix(coverage, value='coverage'):
    # Convert table of coverage into a matrix of coverage at x site
    return pd.pivot_table(coverage, values=value, index='mag', columns='station', aggfunc=np.sum)

def compare_coverage(bins, value='coverage'):
    # bins should be a list of bin names to look at
    if bins is None:
        bins = list(coverage['mag'])
    if len(bins) == 0:
        bins= list(coverage['mag'])
    cov_mat = coverage_matrix(coverage, value)
    mag_frame = cov_mat[cov_mat.index.isin(bins)]
    fig, ax = plt.subplots(figsize=(10,6), dpi=150)
    lines = ax.plot(mag_frame.T)
    ax.tick_params(axis='both', which='major', labelsize=8)
    ax.tick_params(axis='both', which='minor', labelsize=8)
    plt.legend(lines, list(mag_frame.index.values))
    return None
```

In [3]:

```
# Define data location
PRO_COV_FILE = 'merged.csv'
STATIONS_FILE = 'stations.csv'
STATION_ORDER = ['ARK_118M', 'ARK_135M', 'ARK_20M', 'ARK_20M.1', 'ARK_7M', 'ARK_5M', 'ARK_3M', 'ANT_2', 'ANT_4', 
                 'ANT_8', 'ANT_10', 'ANT_15']
STATION_ORDER_DICT = dict(zip(STATION_ORDER, range(0,len(STATION_ORDER))))
# ARCTIC_CIRCLE = 66.56608333333332 # Approx location of the Arctic circle, decimal equivalent of 66°33′47.9
DETECTION_THRESHOLD = 0.9

# Load data
coverage = {}
stations = read_stations(STATIONS_FILE)
coverage = read_abundance(PRO_COV_FILE, stations)
# Do renaming
# Sort coverage station
coverage['stat_order'] = coverage.apply(lambda x: STATION_ORDER_DICT[x['station']],axis=1)
coverage = coverage.sort_values('stat_order')
coverage['mag'] = coverage['mag'].map(papername.rename_mag)
coverage['station'] = coverage['station'].map(papername.rename_station)
# list_mags = list(set(coverage['mag']))
# list_mags = sorted(list_mags, key = lambda x: (STATION_ORDER_DICT[x[:-2]], x[-1]))
```

In [4]:

```
def lowest_class(lineage):
    # Find the lowest classification from a CheckM lineage string
    if isinstance(lineage, float):
        return 'Unknown'
    if len(lineage.strip()) == 0:
        return 'Unknown'
    linpart = [x.strip() for x in lineage.split(';') if x.strip() != 'unclassified']
    return linpart[-1]

def split_heatmap(data, norm_dict, tax_dict, hspace=0.42):
    # Attempt to make a cleaner heatmap version using plotly
    # Convert into pivot table like format
    piv = data.copy().pivot(index='mag', columns='station', values='coverage')

    # Sort by taxonomy
    piv['tax'] = piv.index.map(lambda x: tax_dict[x])
    piv = piv.sort_values(by=['tax'])
    piv = piv.drop(columns=['tax'])

    # Add lowest classfication to index
    piv.index = piv.index.map(lambda x: f'{x} [{lowest_class(tax_dict[x])}]')

    # Normalise counts
    for station, gbp in norm_dict.items():
        piv[station] = piv[station].map(lambda x: x / norm_dict[station])

    # Need min and max values for coverage
    cov_min = piv.min().min()
    cov_max = piv.max().max()

    # Split into Polar and Non-Polar, and plot side by side
    sections = [('Polar', piv.loc[[x for x in piv.index if x[0] == 'P']]), 
                ('Non-Polar', piv.loc[[x for x in piv.index if x[0] == 'N']])]

    # Do some plot setup
    # For each heatmap, want to place two heatmaps close together
    # One using blue-scale (for polar) and one with red-scale (for non-polar)
    barspace = 0.1
    avail = 1 - (hspace * (len(sections)-1)) - barspace
    availper = avail / len(sections)
    width = (2 * len(sections)) + len(sections)
    cw = ([
        (len([x for x in piv.columns if x[0] == 'P']) / len(piv.columns)) * availper,
        (len([x for x in piv.columns if x[0] == 'N']) / len(piv.columns)) * availper,
        hspace
    ] * len(sections))[:-1] + [barspace]
    fig = make_subplots(
        rows = 1, 
        cols = width, 
        column_widths = cw, 
        vertical_spacing = 0,
        horizontal_spacing = 0,
        specs = [
            [{}] * width
        ]
    )
    # Make a dictionary to determine colors
    station_colors = {
        'Polar': px.colors.sequential.Blues[2:],
        'Non-Polar': px.colors.sequential.Reds[2:]
    }

    # Draw the plots
    i = 1
    showscale = True
    for name, data in sections:
        first = True
        # Split the data into polar and non-polar stations
        labelled = data.copy()
        stat_sects = [('Polar', labelled.loc[:, [x for x in list(labelled.columns) if x[0] == 'P']]),
                      ('Non-Polar', labelled.loc[:, [x for x in list(labelled.columns) if x[0] == 'N']])]
        for statname, statdata in stat_sects:
            # Stick on a fake value to force display if all NaN
            statdata['badstat'] = [0] * len(statdata.index)
            color = station_colors[statname]
            fig.add_trace(
                go.Heatmap(
                    z = statdata.values, 
                    x = list(statdata.columns), 
                    y = list(statdata.index),
                    colorscale = color,
                    zmin = cov_min,
                    zmax = cov_max,
                    showscale = showscale,
                    colorbar = dict(
                        thickness = 20,
                        x = 0.86 + (0.07 * i)
                    )
                ), 
                row=1, col=i
            )
            fixrange = [-0.49, len(statdata.columns)-2+0.49]
            fig.update_xaxes(
                dict(tickmode='linear', range=fixrange),
                row=1, col=i
            )
            fig.update_yaxes(
                dict(tickmode='linear'),
                row=1, col=i
            )
            if not first:
                # Hide text on axes
                fig.update_yaxes(
                    dict(showticklabels=False),
                    row=1,
                    col=i
                )
            first = False
            i += 1
        i += 1
        showscale = False
    return fig
```

In [5]:

```
# Plot for prokaryotes
import plotly.graph_objects as go
from plotly.subplots import make_subplots
import plotly.express as px
import plotly
import math

# Set the orca path
plotly.io.orca.config.executable = '/home/hal/miniconda3/bin/orca'

# Normalise count to coverage per gbp
norm_col = 'reads'
norm_div = 1000000
reads = pd.read_excel('summary_statistics.xlsx', sheet_name='bac_summary')[['station_paper', norm_col]].dropna()
pro_gbp_dict = dict(zip(
    reads['station_paper'],
    reads[norm_col] / norm_div
))

# Attach a taxonomy, and sort by this
tax = pd.read_excel('summary_statistics.xlsx', sheet_name='bacbinned').iloc[0:122]
tax_dict = dict(zip(
    tax['MAG ID'].map(lambda x: x[:-1]),
    tax['Bin Lineage']
))

det = coverage[coverage['detection'] > DETECTION_THRESHOLD]

fig = split_heatmap(det, pro_gbp_dict, tax_dict)
fig.update_layout(height=1100, width=1000, title_text="Coverage per millions reads for prokaryotic MAGs")
fig.show()

# Set up local orca install
plotly.io.orca.config.executable = '/Users/duncana/anaconda3/envs/assembly/bin/orca'
#fig.write_image('fig_procoverage.svg')
```

In [6]:

```
# Plot for Eukaryotes
# Define data location
EUK_COV_FILE = 'euk_coverage.csv'

# Load data
euk_coverage = {}
euk_coverage = read_abundance(EUK_COV_FILE, stations)

# Do renaming
# Sort coverage station
euk_coverage['stat_order'] = euk_coverage.apply(lambda x: STATION_ORDER_DICT[x['station']],axis=1)
euk_coverage = euk_coverage.sort_values('stat_order')
euk_coverage['mag'] = euk_coverage['mag'].map(papername.rename_mag)
euk_coverage['station'] = euk_coverage['station'].map(papername.rename_station)

# Normalise count to coverage per gbp
norm_col = 'reads'
norm_div = 1000000
reads = pd.read_excel('summary_statistics.xlsx', sheet_name='euk_summary')[['station_paper', norm_col]].dropna()
euk_gbp_dict = dict(zip(
    reads['station_paper'],
    reads[norm_col] / norm_div
))

# Attach a taxonomy, and sort by this
#tax = pd.read_excel('summary_statistics.xlsx', sheet_name='bacbinned').iloc[0:122]
# ttax = dict(zip(set(euk_coverage['mag']), ['Eukaryote'] * len(set(euk_coverage['mag']))))
# tax_dict = dict(zip(~
#     tax['MAG ID'].map(lambda x: x[:-1]),
#     tax['Bin Lineage']
# ))
euk_tax = {
    'NP3_1': 'Eukaryota; Viridiplantae; Chlorophyta; Mamiellophyceae; Mamiellales; Bathycoccaceae; Bathycoccus',
    'NP2_2': 'Eukaryota; Viridiplantae; Chlorophyta; Mamiellophyceae; Mamiellales; Bathycoccaceae; Bathycoccus',
    'NP2_1': 'Eukaryota; Viridiplantae; Chlorophyta; Mamiellophyceae; Mamiellales; Bathycoccaceae; Ostreococcus', 
    'P3a_3': 'Eukaryota; Viridiplantae; Chlorophyta; Mamiellophyceae; Mamiellales; Mamiellaceae 1; Micromonas', 
    'P5_1': 'Eukaryota; Viridiplantae; Chlorophyta; Mamiellophyceae; Mamiellales; Mamiellaceae 2; Micromonas', 
    'P6_3': 'Eukaryota; Viridiplantae; Chlorophyta; Mamiellophyceae; Mamiellales; Mamiellaceae 3; Micromonas', 
    'P2_1': 'Eukaryota; Viridiplantae; Chlorophyta; Mamiellophyceae; Mamiellales; Mamiellaceae 4; Micromonas',
    'P6_2': 'Eukaryota; Sar; Stramenopiles; Ochrophyta 1; Bacilliarophyta',
    'P6_1': 'Eukaryota; Sar; Stramenopiles; Ochrophyta 2; Bacilliarophyta',
    'P1_1': 'Eukaryota; Sar; Stramenopiles; Ochrophyta 3; Bacilliarophyta',
    'P1_2': 'Eukaryota; Sar; Stramenopiles; Ochrophyta 4; Bacilliarophyta',
    'P3a_2': 'Eukaryota; Sar; Stramenopiles; Ochrophyta 5; Bacilliarophyta',
    'P3a_4': 'Eukaryota; Sar; Stramenopiles; Ochrophyta 6; Bacilliarophyta',
    'NP5_1': 'Eukaryota; Sar; Stramenopiles; Ochrophyta 7; Bacilliarophyta',
    'P2_3': 'Eukaryota; Sar; Stramenopiles; Ochrophyta 8; Bacilliarophyta',
    'P2_2': 'Eukaryota',
    'P1_3': 'Eukaryota',
    'P3a_1': 'Eukaryota; Haptista; Haptophyta'
    
}

euk_det = euk_coverage[euk_coverage['detection'] > DETECTION_THRESHOLD]

fig = split_heatmap(euk_det, euk_gbp_dict, euk_tax, 0.2)
fig.update_layout(title_text="Coverage per million reads for eukaryotic MAGs", width=1000, height=400)
fig.show()
#fig.write_image('fig_eukcoverage.svg')
```

### Percentage of MAGs shared by adjacent stations¶

In [37]:

```
# Make sets of MAGs which are present in each station (regardless of coverage)
piv = det.copy().pivot(index='mag', columns='station', values='coverage')
epiv = euk_det.copy().pivot(index='mag', columns='station', values='coverage')
present = [set(piv[piv[x] > 0].index) | set(epiv[epiv[x] > 0].index) for x in piv.columns]
pc_shared = []
for i in range(0, len(present)-1):
    l = present[i]
    r = present[i+1]
    shared = l & r
    pc_shared.append(len(shared) / len(l | r))
from scipy import stats
print(pc_shared)
print(stats.describe(pc_shared))
# 2nd highest
print(stats.describe([x for x in pc_shared if x < 1]))
```

```
[0.40476190476190477, 0.3333333333333333, 0.6341463414634146, 0.2926829268292683, 0.07246376811594203, 0.7017543859649122, 0.5892857142857143, 1.0, 0.44, 0.896551724137931, 0.7222222222222222]
DescribeResult(nobs=11, minmax=(0.07246376811594203, 1.0), mean=0.5533820291922402, variance=0.07581776174423191, skewness=-0.031095568519956405, kurtosis=-0.7781987910434194)
DescribeResult(nobs=10, minmax=(0.07246376811594203, 0.896551724137931), mean=0.5087202321114643, variance=0.05986258271211491, skewness=-0.17750838906517072, kurtosis=-0.7308646873047269)
```

### Summary of missing results¶

In [7]:

```
piv = coverage.pivot(index='mag', columns='portal', values='coverage')
nans = piv.loc[:,list(piv.isna().any())]
nans = nans[nans.isnull().any(1)]
for f in nans.iterrows():
    series = f[1]
    full = set(series.index)
    nonnan = set(series.dropna().index)
    missing = full - nonnan
    for miss in missing:
        print(f[0], miss)
```

#### Eukaryote / prokaryote correlations¶

In [8]:

```
CORR_THRESHOLD = 0.94
# Pivot data
euk_tbl = euk_det.pivot(index='mag', columns='station', values='coverage')
pro_tbl = det.pivot(index='mag', columns='station', values='coverage')

# Normalise counts & attach taxonomy
for station, gbp in pro_gbp_dict.items():
    pro_tbl[station] = pro_tbl[station].map(lambda x: x / gbp)
# Sort by taxonomy
pro_tbl['tax'] = pro_tbl.index.map(lambda x: tax_dict[x])
pro_tbl = pro_tbl.sort_values(by=['tax'])
pro_tbl = pro_tbl.drop(columns=['tax'])
# Add lowest classfication to index
pro_tbl.index = pro_tbl.index.map(lambda x: f'{x} [{lowest_class(tax_dict[x])}]')

for station, gbp in euk_gbp_dict.items():
    euk_tbl[station] = euk_tbl[station].map(lambda x: x / gbp)
# Sort by taxonomy
euk_tbl['tax'] = euk_tbl.index.map(lambda x: euk_tax[x])
euk_tbl = euk_tbl.sort_values(by=['tax'])
euk_tbl = euk_tbl.drop(columns=['tax'])
# Add lowest classfication to index
euk_tbl.index = euk_tbl.index.map(lambda x: f'{x} [{lowest_class(euk_tax[x])}]')
    
# Merge the pro and euk data
# Add some suffixes
def add_suff(x, suff):
    parts = x.split('[')
    return parts[0][:-1] + suff + ' [' + parts[1]
    
euk_tbl.index = euk_tbl.index.map(lambda x: add_suff(x, 'E'))
pro_tbl.index = pro_tbl.index.map(lambda x: add_suff(x, 'P'))
all_tbl = euk_tbl.append(pro_tbl).fillna(0)

# Make correlations
corr = all_tbl.T.corr()

# Restrict one axis to euk, one to pro
corr = corr[[x for x in corr.columns if 'E [' in x]]
corr = corr.loc[[x for x in corr.index if 'P [' in x]]

# Restrict to only those with corr >= threshold
rmax = corr.max(axis=1).map(abs)
rmax = rmax[rmax > CORR_THRESHOLD]
cmax = corr.max().map(abs)
cmax = cmax[cmax > CORR_THRESHOLD]
corr = corr.loc[list(rmax[rmax > CORR_THRESHOLD].index)]
corr = corr[cmax.index]

corrplot = go.Figure(
    data = go.Heatmap(
        z=corr.values, x=corr.columns, y=corr.index
    )
)
corrplot = corrplot.update_layout(
    height=1800, width=800, 
    title='Correlation between normalised covered of eukaryotes / prokaryotes, ',
    xaxis=dict(nticks=len(corr.index))
)
# corrplot.show()
```

Below is a heatmap showing the Pearson correlation between the normalised coverage pairs of eukaryote & prokaryotic MAGs.
The coverage has been normalised to coverage per million reads.
Only MAGs where some pair has a correlation greater than 0.8 are shown.

Quite a lot of the high correlations are from cases where there are many stations where coverage is (0,0), with one or two stations where coverage is high. Scatter plots for the data are shown further down.

In [9]:

```
corrplot.show()
```

##### Scatter plots for pair with high correlation¶

Scatter plots showing the data for pairs with a correlation greater than 0.8.
Each point represents coverage at in a sample.
Many of them don't have a convincing correlation, with lots of points at (0,0) then one or two stations where coverage is high.

Probably them most interesting are correlations between NP2\_1E (Ostreococcus) and NP5\_3P, NP4\_18P (both Alteromonas), which has a pretty linear relationship. There is one station where the Alteromonas aren't present with higher Ostreococcus coverage.

There could be some associat between P5\_1E, P3a\_3E (both Micromonas) and P5\_24P (SAR86 cluster bacterium)?

In [10]:

```
import plotly.express as px
# Define minimum number of non-zero points must be included for this to be of interest
MIN_POINTS = 3
if MIN_POINTS is None:
    MIN_POINTS = 0
# Loop through eukaryotes
plot_count = 0
plot_data = {}
for euk_mag in corr.columns:
    # Select prokaryotes which have high correlation
    mag_corr = corr[euk_mag]
    mag_corr = mag_corr[abs(mag_corr) > CORR_THRESHOLD]
    filt_tbl = all_tbl.loc[[euk_mag]].stack().to_frame()
    filt_tbl.columns = [euk_mag]
    euk_cov = all_tbl.loc[euk_mag]
    euk_cov = euk_cov.to_frame(name='cov_euk')
    pro_cov = all_tbl.loc[mag_corr.index]
    pro_cov = pro_cov.T.stack().to_frame(name='cov_pro').reset_index()
    pro_cov = pro_cov.merge(euk_cov, how='outer', left_on=['station'], right_on=['station']).fillna(0)
    non_zero = pro_cov[['cov_euk', 'cov_pro']].sum(axis=1)
    non_zero = non_zero[non_zero > 0]
    nonzero_cov = pro_cov.loc[non_zero.index]
    # Make a table of how many non-zero a mag has
    nz_count = nonzero_cov.groupby('mag').count()
    gtr = nz_count[nz_count['station'] >= MIN_POINTS]
    pro_cov = pro_cov[pro_cov['mag'].isin(gtr.index)]
    min_x, max_x = pro_cov['cov_euk'].min(), pro_cov['cov_euk'].max()
    x_margin = (max_x - min_x) * 0.05
    min_y, max_y = pro_cov['cov_pro'].min(), pro_cov['cov_pro'].max()
    y_margin = (max_y - min_y) * 0.05
    if len(gtr) == 0:
        continue
    plot_count += len(gtr)
    plot_data[euk_mag] = pro_cov
    
# Limit this to specific eukaryotes
LIMIT = ['NP2_1E', 'P3a_3E', 'P5_1E', 'P6_3E']
limit_data = {}
for mag, data in plot_data.items():
    if mag.split(' ')[0] in LIMIT:
        limit_data[mag] = data
plot_data = limit_data


# Plot this all on a single figures with multiple subplots
cols = 4
rows = len(plot_data)
fig = make_subplots(rows=rows, cols=cols, horizontal_spacing=0.075)
row = 1
col = 1
colors = {}
colors['P'] = list(px.colors.sequential.Blues)
colors['P'] = list(reversed(colors['P']))[0::1]
colors['N'] = list(px.colors.sequential.Reds)
colors['N'] = list(reversed(colors['N']))[0::
                                    1]
for euk_mag, data in plot_data.items():
    # Select a color
    sel_color = colors[euk_mag[0]][0]
    colors[euk_mag[0]] = colors[euk_mag[0]][1:]
    pro_mags = set(data['mag'])
    col = 1
    for pro_mag in pro_mags:
        mag_cov = data[data['mag'] == pro_mag]
        fig.add_trace(go.Scatter(
            x=mag_cov['cov_pro'], y=mag_cov['cov_euk'], mode='markers', marker=dict(color=sel_color)),
            row = row,
            col = col
         )
        ttext = euk_mag if first or col == 1 else ''
        fig.update_xaxes(range=[min_x - x_margin, max_x + x_margin], row=row, col=col, title_text=pro_mag)
        fig.update_yaxes(range=[min_y - y_margin, max_y + y_margin], row=row, col=col, title_text=ttext)
        col += 1
    row += 1
#     def retitle(x):
#         promag = x.text.split("=")[-1]
#         procorr = mag_corr[promag]
#         text = promag + '<br>r=' + str(round(procorr, 2))
#         x.update(text=text)
# fig.for_each_annotation(retitle)
fig.update_layout(font=dict(size=8), width=800, height=800, showlegend=False)
fig.show()
plotly.io.orca.config.executable = '/home/hal/miniconda3/bin/orca'
fig.write_image('fig5_association.svg')
```

```
---------------------------------------------------------------------------
NameError                                 Traceback (most recent call last)
<ipython-input-10-6c8d9e2265aa> in <module>
     68             col = col
     69          )
---> 70         ttext = euk_mag if first or col == 1 else ''
     71         fig.update_xaxes(range=[min_x - x_margin, max_x + x_margin], row=row, col=col, title_text=pro_mag)
     72         fig.update_yaxes(range=[min_y - y_margin, max_y + y_margin], row=row, col=col, title_text=ttext)

NameError: name 'first' is not defined
```

In [ ]:
