## Supplementary material for "Metagenome-assembled genomes of phytoplankton communities across the Arctic Circle": Suppl. Data 5

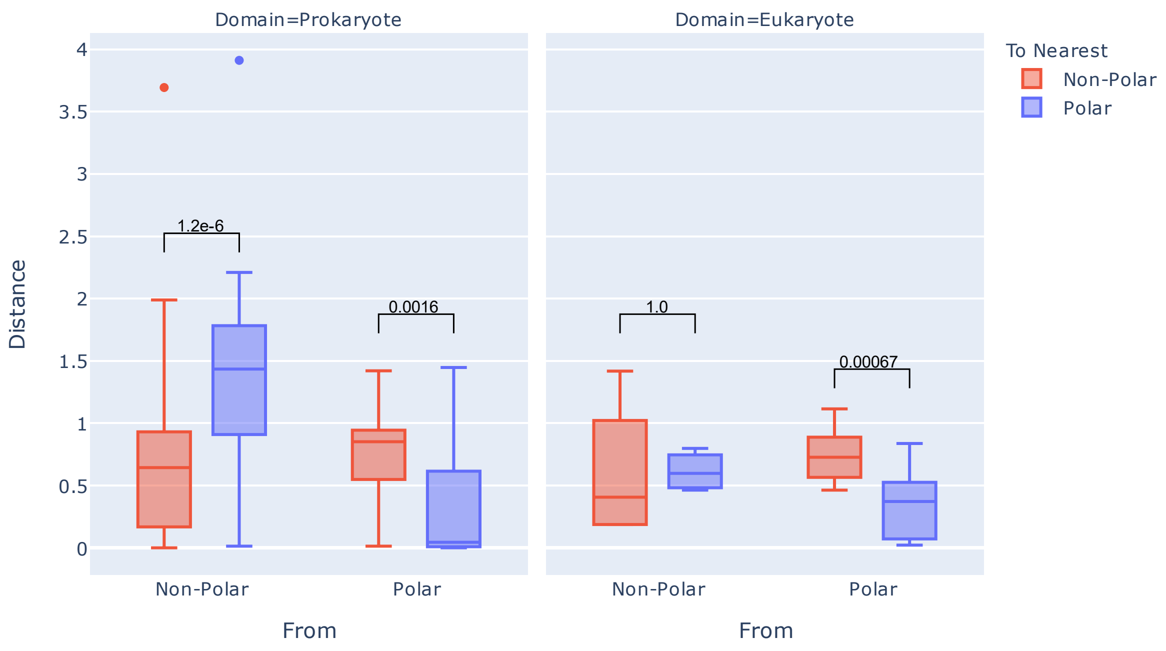


Supplementary Data 5

This figure shows tree distances from MAGs to the nearest polar or non-polar MAG. Values between pairs are p-values from Mood's median test.
