## Supplementary figures and images for "Metagenome-assembled genomes of phytoplankton communities across the Arctic Circle"

### alteromonas_ani.png

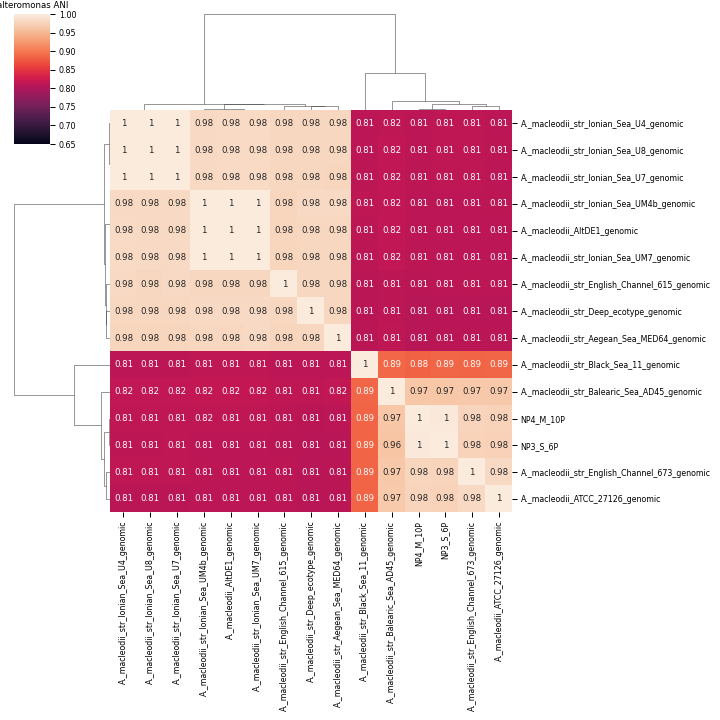

### Bacilliarophyta_ANI.png

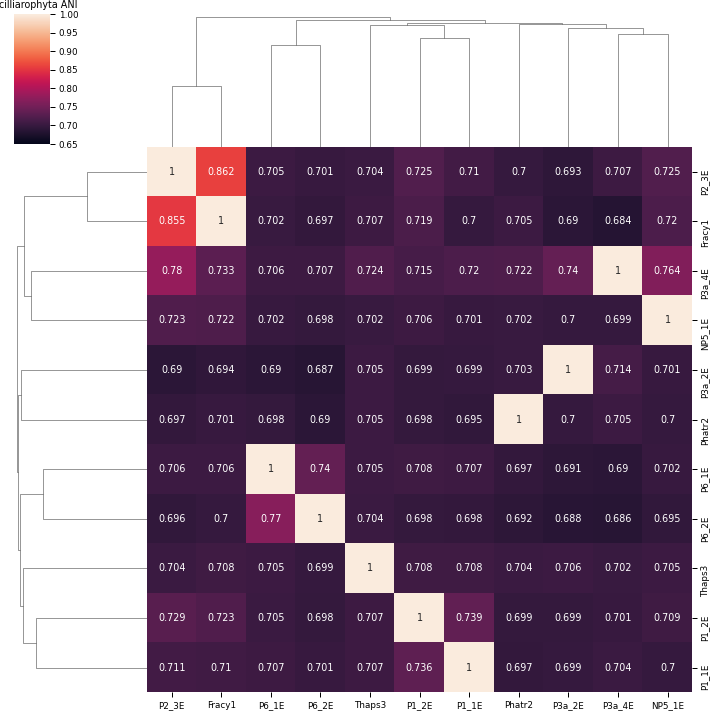

### coraliomargarita_ani.png

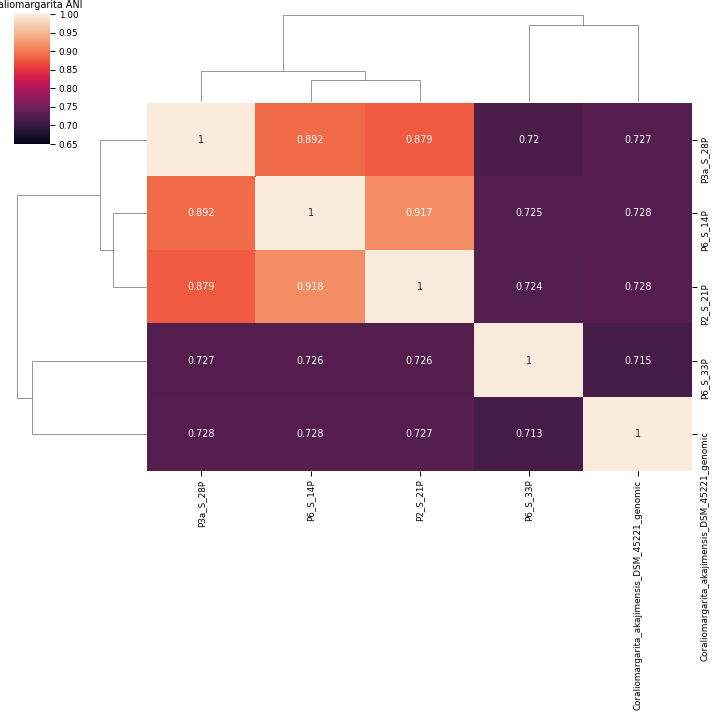

### croceibacter_ani.png

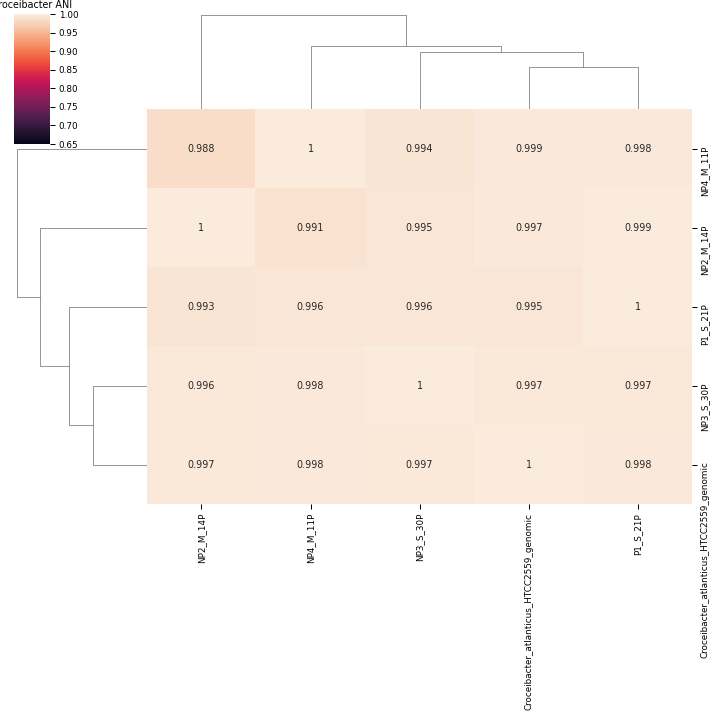

### Haptophyta_ANI.png

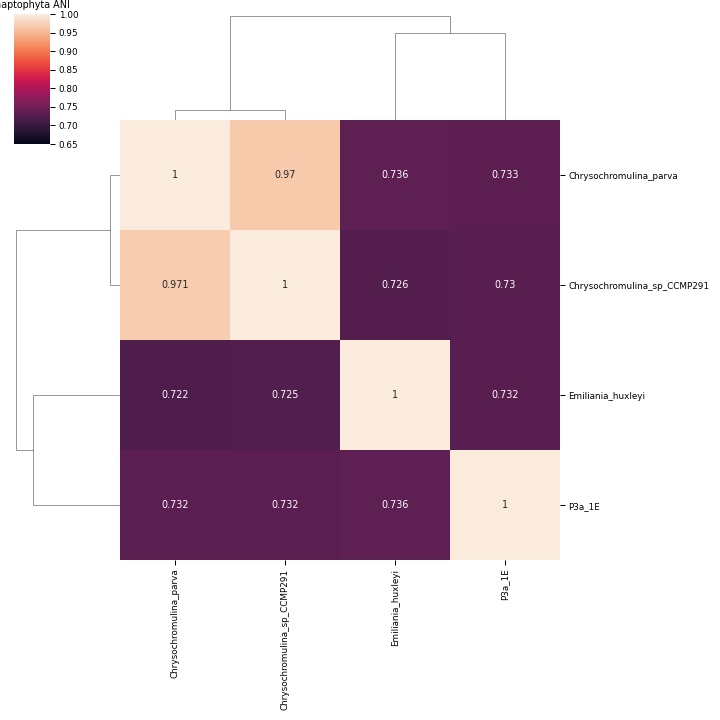

### Micromonas_ANI.png

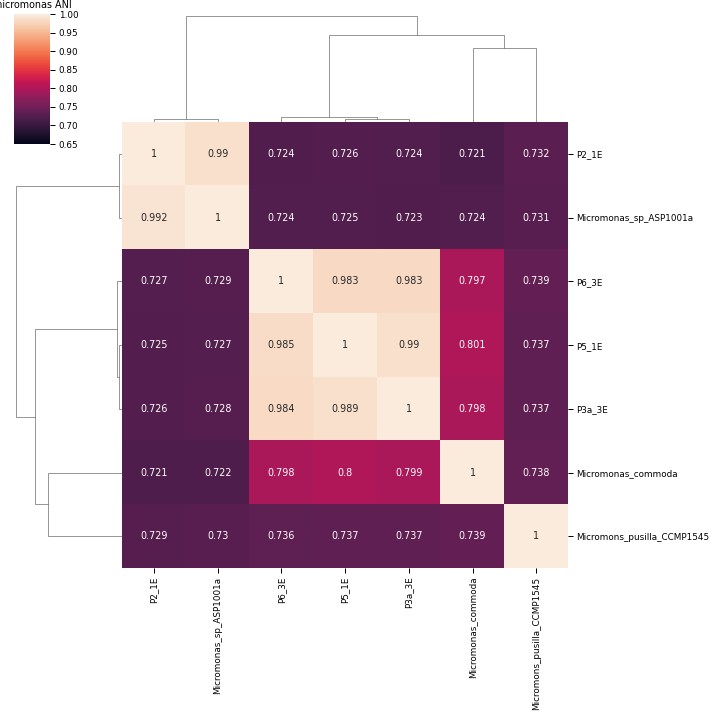

### Suppl. Data 6

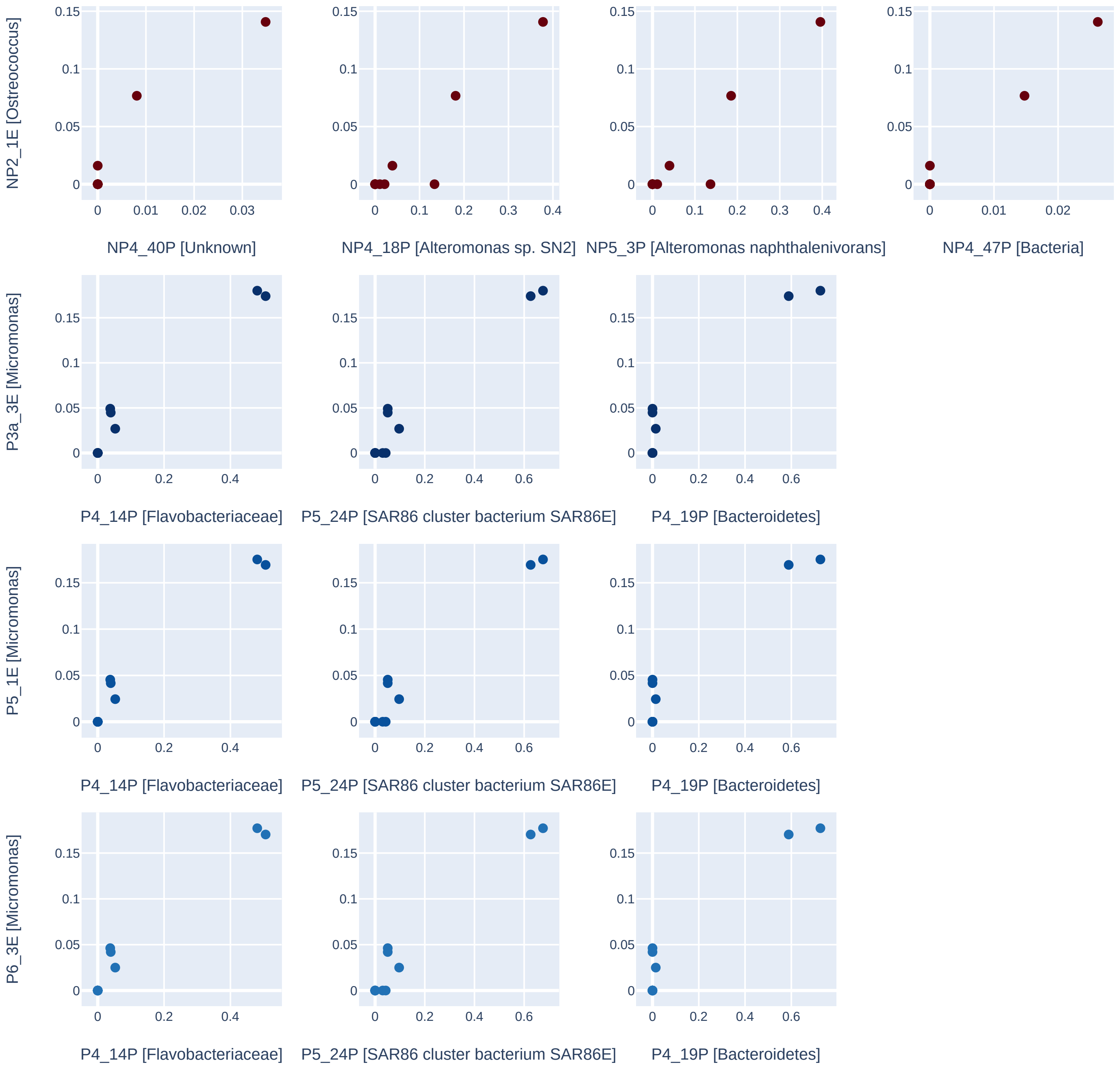
