## Supplementary material for "Metagenome-assembled genomes of phytoplankton communities across the Arctic Circle": Suppl. Data 7: ani_plots.ipynb

```
# %% [code]
import seaborn as sns
import matplotlib.pyplot as plt
import pandas as pd
from glob import glob
import os

# %% [code]
# Function to produce heatmap from a single percentage identity file
def ani_plot(file_loc):
    data = pd.read_csv(file_loc, delimiter='\t', index_col=0)
    # Need to abbreivate the alteromonas names
    if 'alteromonas' in file_loc:
        data.index = data.index.map(lambda x: x.replace('Alteromonas', 'A.'))
        data.columns = data.columns.map(lambda x: x.replace('Alteromonas', 'A.'))
    title = file.split('/')[0]
    title = title.lower().replace('_ani', '')
    h = (5/11) * len(data)
    fs = 0.9 if 'alteromonas' in file_loc else 1
    dp = '.2g' if 'alteromonas' in file_loc else '.3g'
    sns.set_context("paper", font_scale=fs)
    plt.figure(figsize=(h,h), dpi=300)
    sns.clustermap(data, annot=True, fmt=dp, vmin=0.65, vmax=1)
    plt.title(f'{title} ANI')
    plt.savefig(f'{file.split("/")[0]}.png')

# %% [code]
# Loop through percentage identity files and plot
for file in glob('*/*percentage*tab'):
    fig = ani_plot(file)

# %% [code]
```
